# Translatomic database of cortical astroglia across male and female mouse development reveals two distinct developmental phenotypes

**DOI:** 10.1101/681684

**Authors:** Gareth M. Rurak, Stephanie Simard, François Charih, Amanda Van Geel, John Stead, Barbara Woodside, James R. Green, Gianfilippo Coppola, Natalina Salmaso

**Affiliations:** Department of Neuroscience, Carleton University, Ottawa, Ontario; Child Study Center, Yale University; Concordia University, Montreal, Quebec; Department of Systems and Computer Engineering, Carleton University, Ottawa, Ontario

**Author notes:** co-corresponding authors, **Authors for Correspondence**: Natalina Salmaso Carleton University, 1125 Colonel By Drive, Ottawa, Ontario, K1S 5B6, Canada, Gianfilippo Coppola, Yale University, 333 Cedar st, New Haven, CT 06511, USA.

## Abstract

Astroglial cells are emerging as key players in the development and homeostatic maintenance of neurons and neuronal networks. Astroglial cell functions are critical to neuronal migration and maturation, myelination, and synapse dynamics, however little is known about astroglial phenotypic changes over development. Furthermore, astroglial cells express steroid hormone receptors and show rapid responses to hormonal manipulations, however, despite important sex differences in telencephalic regions such as the cortex and hippocampus few studies have examined sex differences in astroglial cells in development and outside of the hypothalamus and amygdala. To phenotype cortical astroglial cells across postnatal development while considering potential sex differences, we used translating ribosome affinity purification together with RNA sequencing (TRAPseq) and immunohistochemistry to phenotype the entire astroglial translatome in males and females at key developmental time points: P1, P4, P7, P14, P35 and in adulthood. Overall, we found two distinct astroglial phenotypes between early (P1-P7) and late development (P14-Adult). We also noted that although astroglia show few basal sex differences in adulthood, they show significant sex differences in developmental gene expression patterns, with peak sex differences observed at P7. At least part of the sex differences observed at P7 appear to be due to males reaching a more mature astroglial phenotype earlier than females. Together, these data clearly delineate and phenotype astroglia across development and identify sex differences in astroglial developmental programs. Importantly, these developmental sex differences could have an impact on the construction and maintenance of neuronal networks and potential developmental windows of vulnerability to neurologic and psychiatric disease.

## Introduction

The cerebral cortex is a complex network of layered neuronal cells that uses a balance of excitatory and inhibitory signals to communicate both within the cerebral cortex itself and to lower brain regions^1^. It is the most recently evolved component of the mammalian central nervous system and is uniquely involved in higher order cognitive functions^2^ that are central to cognitive and emotional processing. The complex layered structure of the cerebral cortex emerges during embryonic development in an “inside-out” pattern consisting of radial glial cell (RGC)-assisted neuroblast migration to lower layers (i.e. VI) before the upper layers are formed ^3, 4^. The appropriate migration of developing neuroblasts is an integral part of the maturation of the cerebral cortex. Early insults or perturbations during this critical period of development may disrupt the gross organization of excitatory and inhibitory networks leading to neurodevelopmental disorders including autism spectrum disorders (ASD), schizophrenia, and mood disorders such as bipolar disorder^5, 6^. Importantly, even subtle modifications in the number or functionality of cortical synapses may lead to serious behavioural consequences and have been related to a number of psychiatric diseases^7^.

In human populations the prevalence of many, if not all, neurodevelopmental disorders and mood disorders related to cortical function is dependent on sex ^8^. Thus understanding sex differences in the organization and function of the cerebral cortex is critical to understanding the neurodevelopmental underpinnings of psychiatric disease. There is a great deal of literature that suggests that gonadal hormones can directly modulate disease states associated with schizophrenia, anxiety and mood disorders^9,10^. Similarly, in rodent models, basal changes in behaviours such as in the forced swim task have been demonstrated across the estrous cycle^11^. Together, these data support an important role for ovarian hormones and in particular, estrogen, in mediating those behaviours related to higher order cognitive functioning in situ. Effects on behaviour and physiology that depend on current levels of hormones such as those described above are referred to as activational. The sex steroid milieu of embryonic and neonatal life, however, can also have organizational effects. These are changes induced in this early stage of development that persist even when the hormonal state producing them has dissipated. Testosterone secreted from the testes of XY-male embryos acting either directly or via aromatisation to estradiol is the primary organizing influence in the sexual differentiation of the brain^12,13^; The role of estradiol in mediating organizational sex differences through neuronal estrogen receptor (ERs) signaling has been extensively studied in the hypothalamus and amygdala. For example, in the medial pre-optic area (MPOA), estrogen receptor signalling induces prostaglandin-mediated synapse formation through microglial and astroglial-mediated functions^13^. Neuroanatomical and functional sex differences are also well defined in various neuronal populations of the hypothalamus^14^, amygdala^15^, and hippocampus^16^. The cerebral cortex, however, has received less attention than other telencephalic regions^17^.

While it is possible that sex differences in cortical network functioning may be completely mediated by direct hormonal modulation, sex differences in the prevalence of anxiety disorders^9^ and ASD^18^ are apparent prior to puberty, and pro-dromal behaviours associated with psychosis also show sex differences early in development^19^, suggesting that organizational sex differences (presumably in cortical networks) may be involved in differences in risk or vulnerability for psychiatric disease.

Astroglial cells have recently emerged as key mediators in brain development and the etiology of neurodevelopmental and mood disorders has been related to astroglial-mediated functions such as neuroblast migration and synapse formation, maturation, and elimination^20,21^. Cortical astroglial cells are a heterogenous cell population traditionally categorised into three main subtypes easily distinguished by morphology, although recent work has suggested that many more subtypes exist when astroglial cells are categorised through single cell RNAseq or other gene and protein expression methods^22,23^. The first subtype of astroglial cells to emerge in development are RGCs^24^. In addition to providing scaffolding for migrating neuroblasts, astroglial cells exhibit stem-like potential in early development and following injury that allows them to give rise to other neuronal cells before maturing and populating the cortex as fibrous or protoplasmic astroglial cells^25,26^. As astroglia mature, their stem cell potential diminishes, as marked by an increase in glutamate synthetase and decreased expression of Sox2 and vimentin^27^. The two main mature morphological subtypes in the cortex are characterized as protoplasmic; canonical astroglia with large domains of many fine processes, and fibrous; astroglial cells with many long fibrous extensions^28^. These mature astroglial cell populations are responsible for the construction and maintenance of both excitatory^29^ and inhibitory^30^ synapses in the developing neocortex, modulating neuron signaling and maturation^31^, and the elimination of synapses during cortical maturation^21^. Work from the Barres laboratory characterised astroglial cells in brain development using next generation sequencing techniques, however these studies did not examine the contributions of biological sex, or early developmental periods^32^. Astroglial cells express ERα^33^, ERβ^34^, and G-protein coupled ERs^35^, and although the full extent of ER functionality in astroglial cells is not understood^36^, astroglia show rapid responses to hormones across various states and manipulations^37^.

Because astroglial cells contribute to cortical development through neuroblast migration, neuronal network integration, and synapse formation, maturation and elimination, and because astroglia show rapid responses to hormones, we hypothesized that cortical astroglial cells may 1) show phenotypic sex differences both overall and across development and 2) execute developmental functions in a sexually dimorphic way. To address this, we employed a transgenic mouse model expressing enhanced green fluorescent protein (eGFP) fused to the L10 ribosome under the pan-astroglial cell promoter AldH-L1 and used immunohistochemistry and TRAPseq to phenotype the astroglial translatome and protein expression. We examined time points pertaining to critical periods of postnatal neocortical development.

## Results

Our initial characterization of neocortical astroglial cells set to measure total numbers of neocortical astroglial cells (astroglial cell number) across development in both males and females. We used unbiased stereology to quantify eGFP+ cells (eGFP was expressed under the control of the pan-astroglial promoter, AldH1l1). No sex differences in total astroglial number were observed (F_1,41_=0.716, p=0.402, Ηη=0.017) (Figure 1A), however there was a main effect of age on this measure (F_5,41_=58.313, p<0.001, Ηη=0.874), whereby the number of astroglial cells plateaued in both males or females after P14 (Figure 1A). Additionally, neocortical volume did not exhibit any significant sexual dimorphism (F_1,41_=0.009, p=0.923, Ηη=0) (Figure 1B). Not surprisingly, we observed a significant increase of brain volume with age ((F_5,41_=135.181, p<0.001, Ηη=0.943) (Figure 1B).

**Figure 1.**
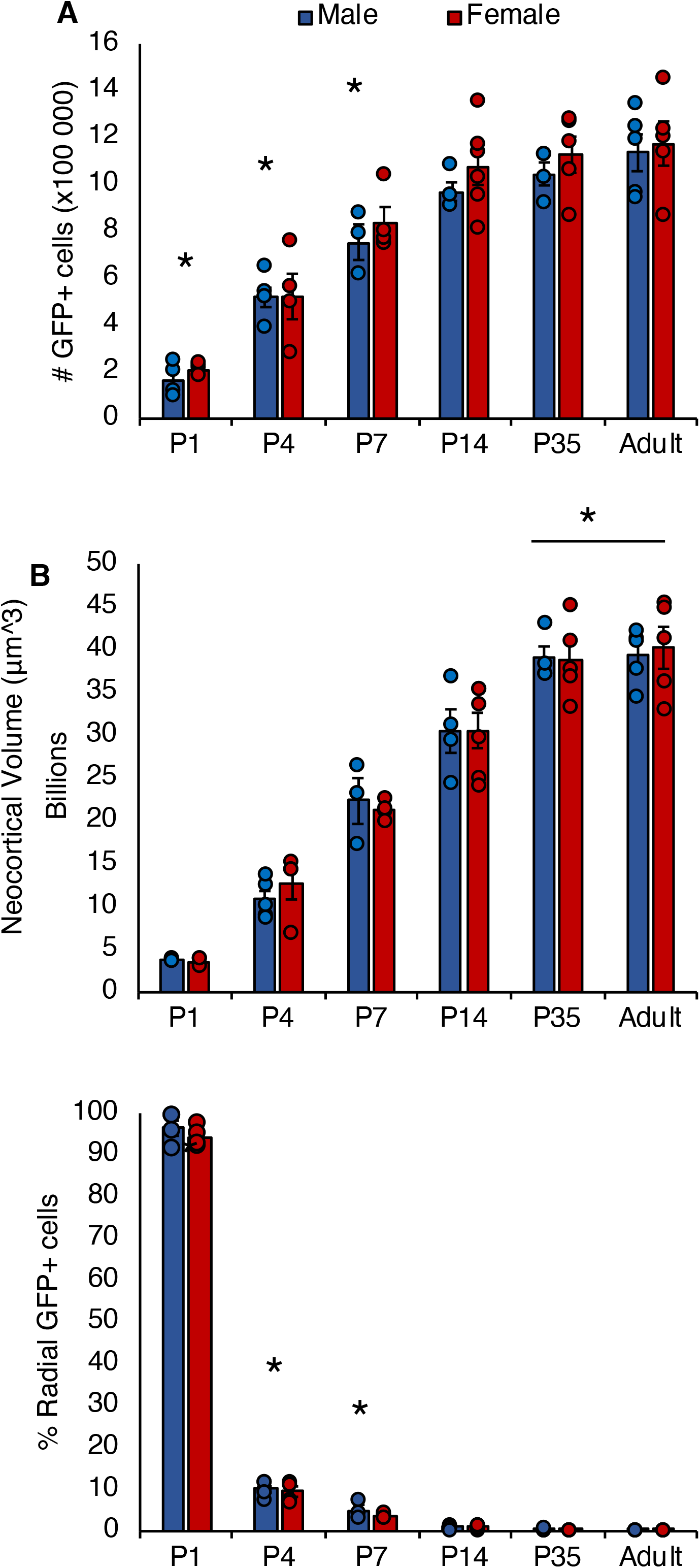
A- Total number of eGFP+ cells in the neocortex. B- Estimated neocortical volume. C- Percentage of radial-like eGFP+ cells. * indicates a significant difference from all other time points. Bars denote group mean and error bars displayed are ± SEM.

Astroglial cells show a great deal of morphological heterogeneity that are typically associated with functionality (Sofroniew & Vinters, 2010). One difference in morphology that is seen during cortical development is an early abundance of the radial phenotype that is replaced by protoplasmic astroglia over development. We assessed gross morphological changes in development (from radial to non-radial) across age to ascertain whether sex differences exist. eGFP+ cells were characterized as radial or non-radial based on the presence of a leading, radial process in the neocortex. No significant sex effects were found (F_1,41_=0.004, p=0.953, Ηη=0), however there was a significant effect of age (F_5,41_=89.291, p<0.001, Ηη=0.916) (Figure 1C). Nearly all (∼95% in both males and females) astroglial cells in the neocortex at P1 exhibited radial morphology (Figure 1C).

To generate a more complete characterization of neocortical astroglia over development and in relation to sex, we employed TRAPseq to assess the astroglial-specific translatome in males and females at each time point. Gene expression levels (as RPKM) for all expressed genes can be found in our searchable database at http://cubic.ca/stardb (see Methods). To validate our TRAPseq methodology, we used q-RT-PCR to compare the unbound fragment to its bound counterpart on levels of two astroglial genes (*GFAP and Glul)* and two neuronal genes (*Rbfox3 and Pvalb*). As expected, we found a large increase in astroglial gene expression in the bound fragment and a large reduction in neuronal-specific gene expression, suggesting a successful enrichment of the astroglial translatome (Supplemental Figure 3). In order to further characterise astroglial-specific markers by age and sex, we plotted RPKM data for each of the top 20 genes expressed only in postnatal neocortical astroglia from both the Cahoy dataset^32^, 28 genes from the Zhang dataset^38^ and 48 genes from the Rowitch dataset that identified layer-specific astroglial cell gene expression (Bayaktar et al. 2020) that were non-overlapping (Supplemental Figures 4, 5 & 6). Interestingly, expression of most astroglia-specific genes from the Zhang and Cahoy datasets (27/48) increases across development, for example *CYP4F14*, a Vitamin E hydoroxylase and *SLC39A12*, a zinc transporter involved in the pathophysiology of schizophrenia^39^ both have low expression levels early on and rise to adult levels at P14 (Supplemental Figure 4). Only one astroglial gene decreased in expression over development, *Pla2g3,* a gene involved in oxidative stress that has been linked to Alzheimer’s disease^40^.

Supplemental Figure 5 depicts the 14/48 astroglial genes that show stable expression over development and finally, Supplemental Figure 6 depicts those genes (8/48) that show consistent low expression over the entire developmental period assessed. Interestingly, we did not find any apparent sex differences in these astroglial-specific genes, although a few appear as DEGs at specific time points, for example, the cytokine Il33 is a DEG at P7 (lower in females-see below). Supplemental figure 7 shows astroglial cell gene expression for Layers 2-4 and deep layers 5-6, supplemental figure 8 shows upper-layer biased astroglial cell gene expression across development, and supplemental figure supplementary figure 9 shows gene expression for genes identified with the AST2, AST3 subtypes.

### Astroglial cell show age related differences in translatome

Next, we examined the overall sample-to-sample relationship by hierarchical clustering to identify patterns throughout development and across sex. Clustering shows a pattern of organization that is primarily affected by age, with P1, P4 and P7 (early) and P14, P35 and adult mice (late) showing clustered expression patterns that differ from the early time points (Figure 2A). We manually sorted samples by sex and age and although some organization by sex is noted in Figure 2B, this is not as clearly demarcated or as extensive as that observed by age. Indeed, when we collapsed gene expression levels across all time points, only 16 genes were differentially expressed between female and males (Supplementary Data File 2) and a subset of these sex differences in gene expression is for genes are located on the sex chromosomes, suggesting little to no sex differences in neocortical astroglia across all time points overall. However, when we look at sex differences over time, we do note sex differences in developmental patterns. It is clear that there are two major neocortical astroglial phenotypes across age: an early (P1-P7) and late (P14-adult) phenotype. However, male samples show an “intermediate” cluster and begin to show the late or mature astroglial phenotype at P7, whereas females show this phenotype at P14 (Figure 2B).

**Figure 2.**
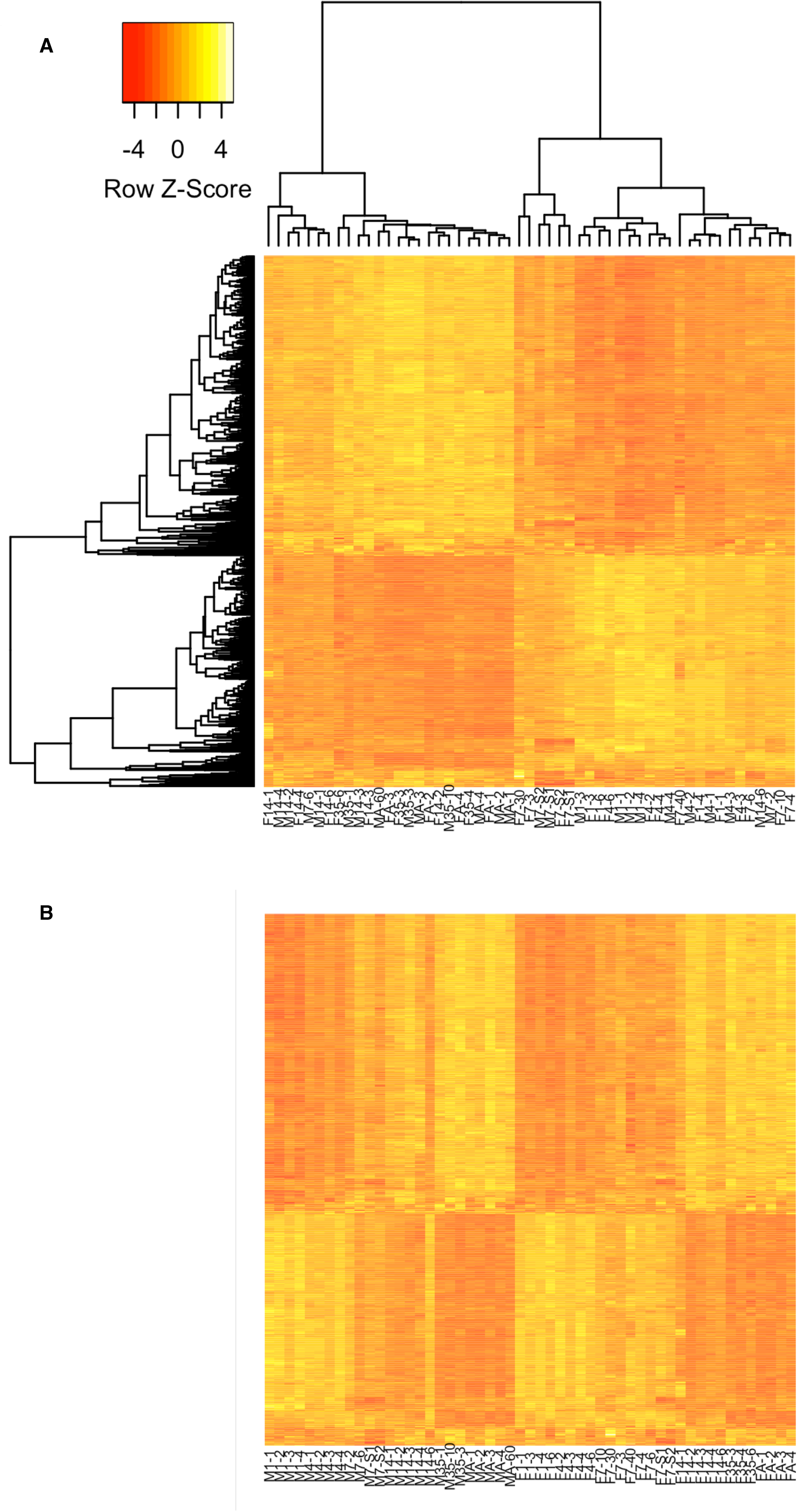
A- Clustered heat map of individual sample gene expression levels from TRAPseq data. B- Represents a heat map of samples manually sorted by sex and age.

To validate the early and late phenotype that we found using TRAPseq, we compared TRAP extracted RNA on six genes differentially expressed between the early and late timepoints, 3 associated with synapse/network maturation (*Gja1, Lynx1, Grin2c*) and three steroid hormone receptors (*Ar, Pgr, Esr1*). Each of these showed a similar pattern of results between TRAPseq and qPCR (Supplemental Figure 10), which is highlighted by the significant positive correlations shown in Supplemental Figure 11. In addition, we further validated the early and late astroglial cell phenotype, by assessing neocortical astroglial cells protein expression using immunohistochemistry of common astroglial markers at similar timepoints. Using the Aldhl1-L10 eGFP+ mice, we quantified expression of the astroglia-specific cytoskeletal filament protein glial acidic fibrillary protein, (GFAP). GFAP expression was not sexually dimorphic across development (F_1,41_=1.259, p=0.268, Ηη=0.029), however, there was a main effect of age (F_5,41_=32.271, p<0.001, Ηη=0.793) (Figure 3A). The number of GFAP+ cells reached peak levels at P14 and remained stable into P35 and adulthood (Figure 3A). Prior to P14, no more than 15% of eGFP+ cells co-expressed GFAP, from P14-onwards approximately 25% of eGFP+ cells expressed GFAP, consistent with previous work that has demonstrated that a large proportion of cortical astroglia do not express GFAP under basal conditions (Simard et al., 2018). Next, we quantified the number of cells that expressed vimentin, another intermediate filament protein in astroglial cells, that is generally associated with a stem-like radial sub-type of astroglial cells. A significant interaction of sex by age was found (F_5,41_=3.871, p=0.006, Ηη=0.321) (Figure 3B,D), such that females showed significantly more vimentin+ cells compared with males at P7 (t=3.41, p=0.011) and males showed significantly more vimentin+ cells in adulthood (t=3.895, p=0.005) (Figure 3B). Because vimentin expression is associated with astroglial cell neurogenic and stem cell potential and because we observed sex differences in vimentin expression, we quantified the proportion of actively dividing eGFP+ cells across the postnatal period. Using a marker of active s-phase cell division, Ki67, we quantified the percent of Ki67+eGFP+ cells. Ki67 was present in less than 2% of the astroglial cell population throughout the postnatal period in both males and females (Figure 3C). There were no significant sex differences across development (F_1,41_=0.862, p=0.358, Ηη=0.021), however there was a main effect of age (F_5,41_=26.799, p<0.001, Ηη=0.766). Importantly, the findings here that the number of astroglial cells expressing vimentin show sex and age differences, and proportion of astroglia expressing GFAP and Ki67 show distinct early and late expression closely resemble the expression patterns revealed by TRAPseq analysis.

**Figure 3.**
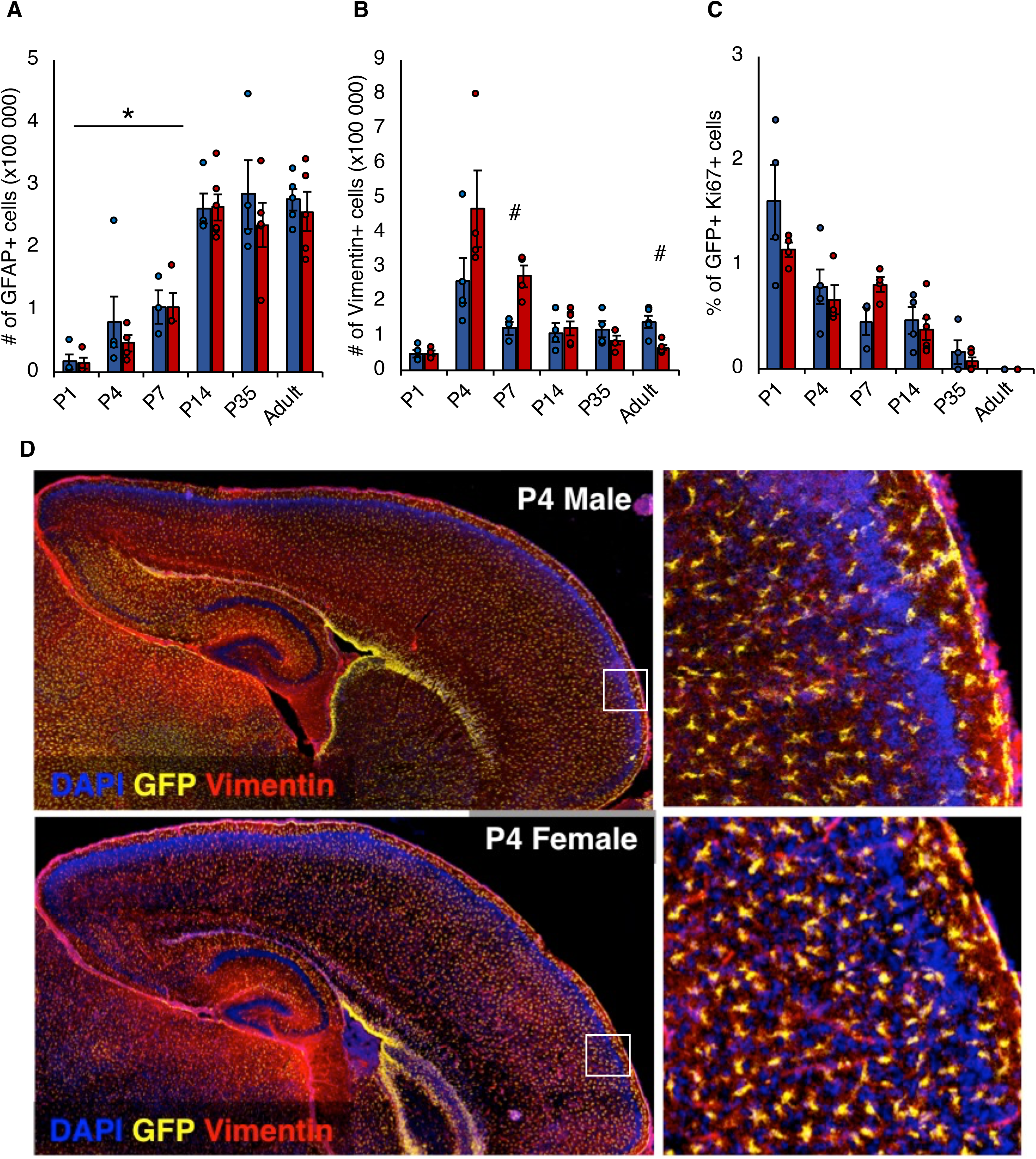
A- Total number of cells expressing GFAP. B- Total number of vimentin+ cells. C- Percent GFP+ co-expressing Ki67. D- Representative mosaic of confocal pictomicrographs at 10x, whole cortex sagittal section (left), zoom on the molecular layer (right). * indicates a significant difference from all other time points; # indicates a significant difference between male and female at that time point. Bars denote group mean and error bars displayed are ± SEM.

### The developmental pattern of the astroglial translatome is sexually dimorphic

To further characterize the *“early”* and *“late”* phenotype observed in the astroglial cell translatome, we examined the number and proportion of up- and downregulated genes in the early versus late developmental phenotypes, both overall and by sex (Figure 4A). The early versus late phenotype in males showed 8983 DEGs whereas the female early versus late phenotype had 9688 DEGs (full list of DEGs in Supplementary Data File 1). Interestingly, the proportion of up- or downregulated genes was similar, with males and females showing 62% and 55% of DEGs in the early versus late phenotype being upregulated, respectively. When we collapsed the DEGs across sex to generate a general early vs late phenotype for astroglial cells, the number of DEGs reaches 12131 with 58% of those genes being upregulated in the late phenotype (Figure 4A). This suggests that the difference in the early and late developmental phenotype is not merely reflective of turning off developmental genes but rather is a combination of downregulating genes associated with early development and turning on genes associated with later neocortical development and basal functioning in early adulthood. Some key DEGs stand out as they have not previously been functionally described in astroglial cells. In particular, *Trim67*, an important and evolutionarily preserved guidance cue for neurite outgrowth^41^, is high in early development and, as expected, decreases with age. Additionally, *Ntng1*, codes for Netrin G1, another guidance cue that is paramount to developing neuroblasts, is higher during the “late” developmental phenotype in astroglial cells. Differential *Ntng1* expression appears when examining both the overall and the sex-specific early versus late astroglial cell phenotype. *Ntng1* has been implicated in the etiology of familial schizophrenia^42^ and *Ntng1* mRNA is in lower cortical samples from individuals with schizophrenia and autism^43^.

**Figure 4.**
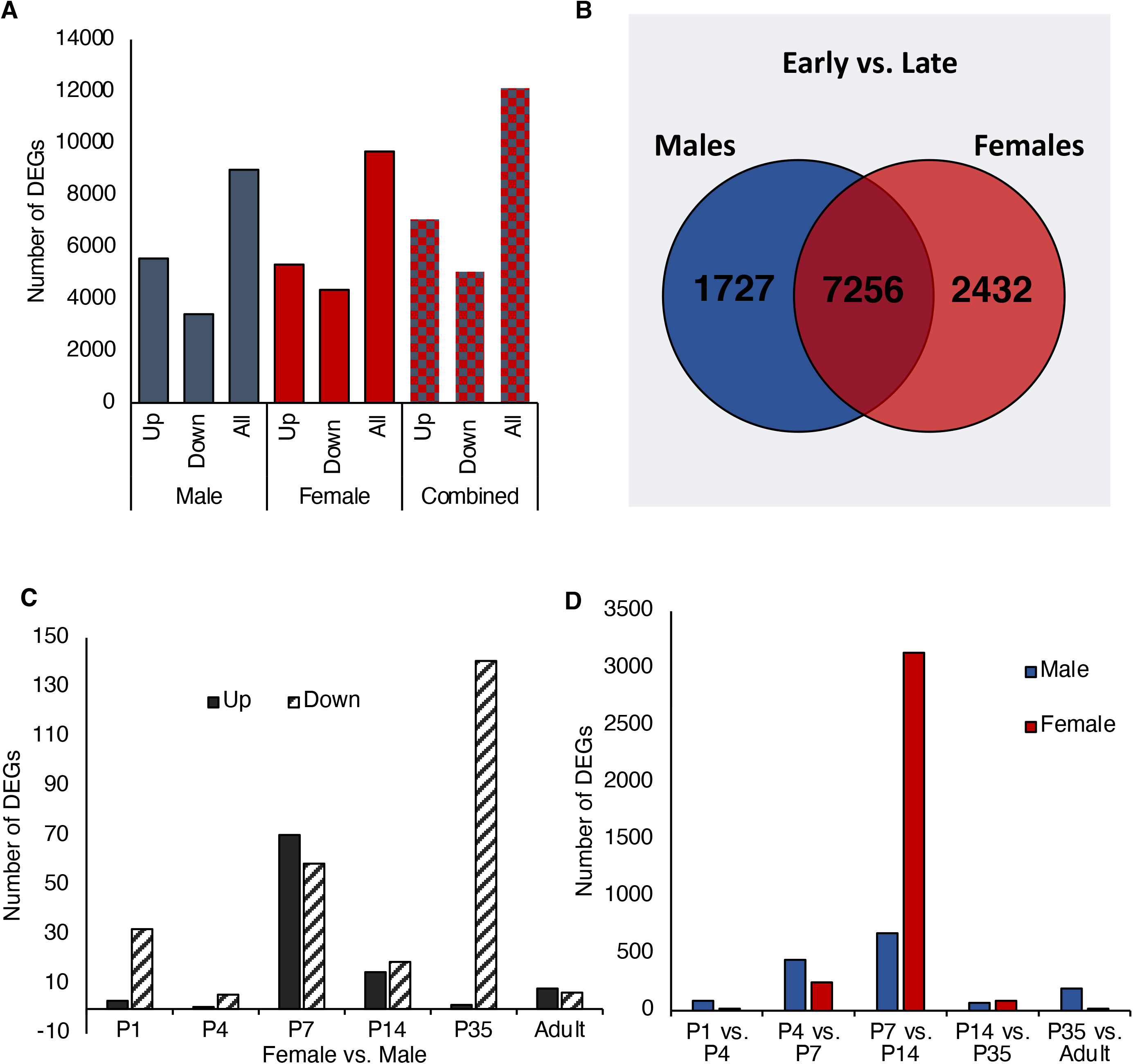
A- Number of DEGs between early and late phenotype in males and females. B- Venn diagram displaying overlap in DEGs between females and males. C- Bar plot showing the number of up- and downregulated DEGs between males and females at each time point (females=reference group). D- Bar plot showing the number of DEGs between time points.

We compared the proportion of overlap between the female-specific and male-specific DEGs in the early vs late phenotype. Of the 11415 unique DEGs, approximately 64% overlap between males and females, however about 21% of total DEGs are specific to females and only about 15% are specific to males (Figure 4B). When we further characterize the sex-specific canonical pathways (CPs) and gene ontology (GO) by gene set over-representation, a proportion of the top five female-specific CPs are related to cell cycle/mitosis, marked by CPs in ribosome function. In comparison, male-specific CPs are related to oxidative phosphorylation and small molecule transport. With respect to GO, a large proportion of the top five female-specific and male-specific GOs are related to intracellular parts and organelles. This suggests that there are distinct developmental patterns in neocortical astroglial cells that emerge between the sexes. To broaden the understanding of sex differences, we identified DEGs of females versus males at each time point (Figure 4C, Supplementary Data File 2). At P1, there are only 20 DEGs, however there are 183 DEGs at P7 and 100 DEGs at P14 (Figure 4C; full list of DEGs in Supplementary Data File 2). Analysis of DEGs at P1 calls one CP related to histone demethylation. When we analyze DEGs between timepoints and sexes, we observe a peak in the number of DEGs when comparing P7 vs 14 in both females and males (Figure 4D and Supplementary Data File 3). Within this peak (P7 vs. P14), 1658 DEGs are identified in males and 3685 DEGs are identified females. These sex differences between P7-P14 are particularly striking given that these are the developmental time points, where sex hormones are largely dormant. This suggests that these sex differences in astroglial gene expression are part of a developmental program induced by perinatal exposure to hormones involved in sexual differentiation and/or directly induced by sex-linked chromosomes. Remarkably 39% of DEGs at P7 and 79% of DEGs at P14 are downregulated in females. Within these downregulated genes are typical markers of cell activity, the immediate early genes *Fos* and *Arc*, suggesting decreased basal activity in female astroglial cells at these timepoints (Supplementary Data File 3, Tab a). Analysis of DEGs at P7 show that three of the top five CPs are related to synapse development and transmission while the remaining two are related to intracellular signalling pathways. Similar analysis of DEGs at P14 show that the majority of the top five CPs are related to cell cycle. In addition, the top five GO terms from DEGs at P7 are related to myelination and synaptic components and at P14, the top five GO terms are related to cell cycle.

A more complete view of sex differences is captured by differential network dynamics. We used co-expression network analysis (see Online Methods), to infer, separately a co-expression network in females and males throughout development, and identified 10 “female” (F) and 12 “male” (M) co-expression modules, plus one unassigned gene module, grey (Supplementary Data File 4 & Supplementary Data File 5, Tab1). In females, the F3, F6, F7 and F9 modules are enriched in genes upregulated in late vs. early, and the F1, F2, F4 and F10 modules are enriched in genes downregulated in late vs. early (see Supplementary Data File 4,Tab4). The F3, F6 and F9 (upregulated) modules are enriched in astroglial genes. The F2 and F4 modules (downregulated) are enriched in cell cycle terms, and the F10 module (downregulated) is enriched in genes involved in proteasome activity and cell cycle. In the males’ network, the M2, M3, M4, M8 and M12 modules are enriched in genes upregulated in late vs. early, and the M1, M7 and M10 modules are enriched in genes downregulated in late vs. early in males (see Supplementary Data File 5, Tab 4). Similar to what was observed in the female network, some of the modules enriched in upregulated genes in the late vs. early phenotype in males (M2, M3, M6 and M8), are enriched in astrocyte genes. It is likely that, in both networks, the astrocyte signature for the modules enriched in upregulated genes reflects a shift in a mature astroglial phenotype during development. Finally, the M10 module (downregulated) is enriched in cell cycle terms, and the M1 module (downregulated) is enriched in genes involved in mRNA processing. The cell cycle signature for some of the downregulated modules likely reflects a reduction of proliferative potential of the cells over time, as would be expected over development.

We conducted an eigengene analysis of the network modules to evaluate how modules changed over development. Overall, we found consistent results across males and females whereby modules enriched with astroglial genes were increased over development, modules enriched with genes involved in cell cycle regulation were downregulated over development, modules involved in translation (ribosomes) were also downregulated and finally modules enriched in genes related to mitochondria and mitochondrial function were increased with age, further lending support to the “early” and “late” astroglial cell phenotypes (Supplementary Data File 5, Tab 4 and Figure 5 A-C).

**Figure 5.**
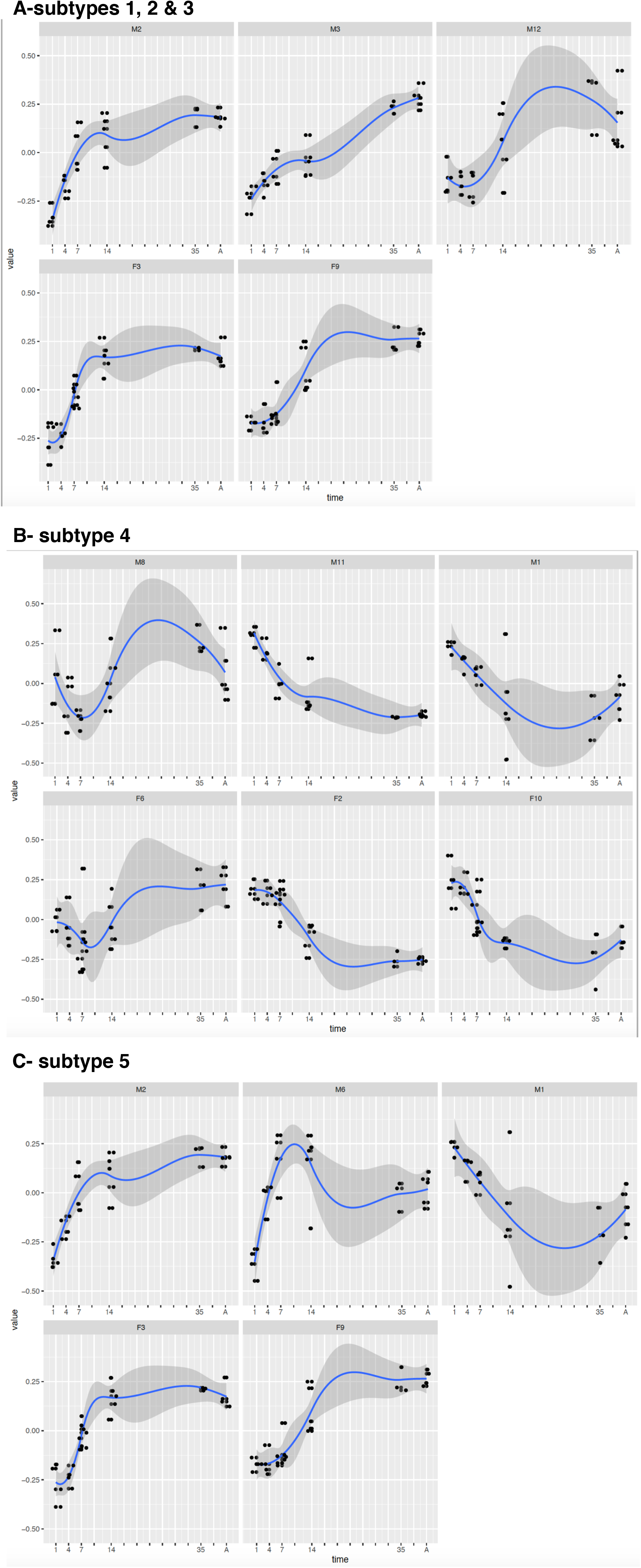
Plot of modules Eigen-value vs time. Represented is the subset of the male and female modules enriched in astrocyte subtype markers. A- modules enriched in AST1, AST2, AST3 subtypes; top row: modules M2, M3, M12; bottom row: F3, F9. B- modules enriched in AST4 subtypes; top row: modules M8, M11, M1; bottom row: F6, F2, F10. C- modules enriched in AST5 subtypes; top row: modules M2, M6, M1; bottom row: F3, F9.

A closer look at the temporal dynamics highlights some interesting features, with the female network characterized, mostly, by monotonic, sigmoid like temporal profiles, and the male network characterized by a larger variety of patterns (subset shown in Figure 5 A-C). In females, the sigmoid profiles are either decreasing with age (F1, F2, F4 and F10) or increasing with age (F3 and F9). We also find two modules (F6 and F7), increasing with age, with a possible peak at P7 and P14, respectively. In the sigmoid profiles, we also identified a ‘slow’ subgroup (F1, F2 and F9), characterized by a slow change between P1 and P7 and more activity around P14; and a ‘fast’ subgroup (F3, F4, F5, F7 and F10), characterized by a fast change around P7 and lower activity from P14 onward. The temporal dynamic pattern analysis is consistent with the “early” and “late” hierarchical clustering. Unlike the female network, most of the modules in the male network show a relatively faster temporal dynamics at P1 to P4-P7. Noticeable are also the pronounced early peaks in the M5 and M6 modules, the late peaks in the M4, M8 and M12 modules and a pronounced deep at P7 in the M10 module. Functional analysis shows that the ‘early peaks modules’, M5 and M6, are enriched in neurotransmitter secretion and synaptic transmission terms, respectively; the ‘late peaks’ modules, M4, M8 and M12, are enriched in transmembrane transport, signaling receptor activity/astrocyte, and transmembrane transport terms, respectively. The M10 module is enriched in mRNA processing, and the M1 in neuron terms.

We tested the modules for enrichment in Estrogen and Testosterone related gene sets (i.e. ESR1 and ESR2 gene targets and genes/proteins interacting with the chemical Testosterone (Rouillard et al., 2016) and found significant overlap between the M2 module in males with the ESR1 targets. The male M2 module expression increases rapidly, from P1 to P14, where it levels off. This may be consistent with the active process of masculinization of the brain early in development. No significant overlap is found with any of the modules in the female set.

### Network dynamics highlight differential astrocyte subtype differentiation

We tested the male and female modules for enrichment in gene marker for astrocyte sub-types (Bayraktar et al., 2020; Nature Neuroscience; Batiuk et al., 2020; Nature Communications) to investigate sexual dimorphism in astrocyte subtype differentiation, and describe them following the developmental timeline (see Supplementary Data File 4, and Supplementary Data File 5, Tab5).

The AST4 (neural stem or progenitor cells) markers overlap with modules F2, F6 and F10 in females, and M1, M8 and M11 in males. The overlap with modules decreasing with time is expected and in line with the stem cell/progenitor phenotype of AST4, while the overlap with F6 (females) and M8 (males) is more surprising, given that both modules first decrease, then increase from P7 (see Figure 5B). Interestingly, vimentin expression is one of the major markers of the AST4 subtype (Batiuk et al., 2020; Nature Communications) (in Supplementary Data File 2), therefore our observation of sex differences in vimentin expression over time (Figure 3B) is consistent with sexual dimorphism in these particular network dynamics.

The AST5 (intermediate progenitors) markers overlap with F3 and F9 in females and M1, M2 and M6 in males (see Figure 5C). All modules show increased expression over time at a faster rate than modules overlapping with the mature astrocytes, and differently in males than in females. Interestingly, in males, we find overlap with module, M1, a module decreasing over time, and lack of overlap with a similar module in females (see Figure 5C). Top scoring terms for this module are mRNA processing, Spliceosome and Proteasome. Down-regulation of this module could be associated with cell cycle and the associated dismantling and ciliogenesis of astroglial cilia (Sterpka and Chen, 2018). Alternatively, the proteasome gene down-regulation could also be consistent with the GFAP increase over time (Middeldorp et al., 2009). The faster temporal dynamics and the male specific module point to sexual dimorphisms in astroglial differentiation dynamics that become explicit at the intermediate progenitor stage; these warrant further investigation in future studies.

The AST1, AST2 and AST3 (mature astrocytes) markers overlap with modules showing increasing expression with time both in females (F9 and F3) and in males (M2, M3 and M12). Notably, the F9 module in females (AST1, AST2) has a slower rise than F3 (AST2, AST3), suggesting a delay in AST3 with respect to AST1 (see Figure 5A). In males, M2 is enriched in all the mature cell types, probably capturing common underlying processes; the M12 (AST1, AST2) module starts increasing late, at P14, M3 has a slower increase with respect to M2, again suggesting a delay in AST3 differentiation with respect to AST1 and AST2. The overlap with modules increasing expression in time is consistent with the mature phenotype of these astroglial classes. Overall, some female modules show little change between P1 and P4, and a steeper increase from P4 to P7, while male modules tend to change from P1, suggesting a delay in females. Also, we find two shared modules in males, across two transitions, M1 in AST4 to AST5, and, M2, in AST5 to AST1/AST2/AST3; conversely, females do not have any shared modules in the first transition, AST4 to AST5, and have two shared modules, F3 and F9, both in the second transition AST5 to AST1/AST2/AST3.

### Differential Transcriptional Regulation Dynamics Drives Sexually Dimorphic Astrocyte Differentiation Programs

The different developmental trajectories and the faster temporal dynamics in males supports the idea of differential regulation of astrocytic differentiation ?between males and females, already present at P1. We then investigated the regulation of network modules to identify determinants of network dynamics and subtype differentiation. We first identified a number of putative transcription factors regulating the male and female network modules and found an average of 320 and 340 significant TFs per module, respectively, for a total of 856 and 858 TFs (see Methods), with 2 male-specific (HELT, KAT7) and 4 female-specific (CARF, CPEB1, MTF1, PHF1 TFs. Interestingly, HELT and CPEB1 target AST4 genes markers involved in different processes (HELT: neural crest differentiation, cell fate specification; CPEB1: cell cycle related; cell projection; cytoskeleton) (see Table GF4 and GF5 Tab6).

We then used the expression of TFs target genes as a reporter of activity, similarly to the Viper algorithm (Alvarez et al., 2016), and estimated TF differential activity between females and males at each time point, and between time points, using the camera algorithm (Wu and Smyth, 2012) (see Methods).

We first identified the subsets of differentially active TFs (DATFs) between P1 and P4 in males and females, as putative determinants of early transcriptional dynamics, resulting, respectively, in a set of 839 and 510 DATFs, respectively. Within this subset, we identified DATFs at P1 between males and females, as putative TFs driving the difference in early transcriptional dynamics, resulting, respectively, in a set of 775 and 119 DATFs, suggesting a stronger activity in transcriptional regulation in males vs females. Next, we focused on the subset of DATFs targeting gene markers of individual astrocyte sub-types, to focus on cell-type specific differential regulation. We found 551 DATFs in males and 33 in females, with targets co-expressed in the modules M1 (57), M11 (494) and F2 (24), F10 (9), and enriched marker genes for AST4. At P1, the already consistent number of differentially active TFs might reflect the differential priming by testosterone and/or aromatization of estradiol in males of astrocyte progenitor cells, which would reach peak levels at this time. Interestingly, ESR1 is a DATFs controlling targets in M11, suggesting that some of the sex differences in the developmental trajectories of cortical astroglia, may be directly initiated by the perinatal exposure to estradiol critical to male sexual differentiation.

We then aggregated the TF-target genes and built a regulome (i.e. a regulatory network represented as a directed graph) for the males (13536 genes and 404928 edges) and females (13756 genes and 449097edges) using known regulatory relationships, and identified the putative TFs, potentially upstream of the other TFs/genes, i.e. the source nodes of the graph (see Methods). The source nodes are the nodes in a directed graph, with a non-zero out-degree and a zero in-degree. We found ESR1, ZFP3, MZF1, PRRX1, SOX7, MSX2, ZGPAT, TCF24, and MYRFL in the male network and BARX2, FOXR2, and MYRFL in the female network. Interestingly, while ESR1 is present in both, it is a source node only in the male network, supporting a central role of ESR1 in AST differentiation in males and of network rewiring early in masculinization.

Early masculinization of the brain occurs largely as a result of a perinatal surge in testosterone that is aromatised into estradiol in the brain of males, therefore we further examined P1 as a critical time point in setting up differential developmental patterns in the astroglia of males and females. Extensive work from the McCarthy laboratory has demonstrated that prostaglandins, and specifically, glial release of cyclooxengase 2 (COX-2) is a key regulator of estrogen-mediated architectural sex differences in the hypothalamus and the amygdala^44,45^. The gene for COX2, *Ptgs2*, is a sex-specific DEG in our data set, in the overall females versus males analysis, in the early female versus male phenotype, and at P1 female versus male comparison (Supplementary Data File 1 & Supplementary Data File 2) Furthermore, Cox2 is listed as an ESR1 target in our dataset and it is well-known to be downstream of ESR1 signalling (Stossi et al., 2004). To validate COX2 expression over development, we counted the number of cortical COX2+ cells in astroglia (co-localised with eGFP) and saw a similar pattern of expression to that observed in our TRAPseq, with a significant sex difference in astroglial Cox2 protein expression at P1 (F_1,41_=12.517 p=0.017, Ηη=0.715)(Figure 6A). COX2-mediated masculinization of the hypothalamus occurs in response to estradiol, which is aromatised from the surge in testosterone in males that occurs perinatally in rodents (Figure 6B). This perinatal surge in testosterone has been linked to sexual differentiation of the hypothalamus, amygdala, and mediates sexually dimorphic juvenile and adult behaviours^46–48^. To validate our findings and to test if astroglial COX2 protein levels are similarly regulated by estrogen in the cortex, we compared vehicle injected males and females with females that were masculinized by injecting estradiol benzoate at P0 and then sacrificed all groups 36 hours later. The overall one-way ANOVA showed a trend for significance (F_3,20_=2.781, p=0.068, Ηη=0.294), however because our previous experiment showed a difference between males and females, we conducted a planned comparison of these groups. Females at P1 show fewer COX2+ cells in the frontal cortex compared to males (F_1,9_=7.827, p=0.021, Ηη=0.465) and females exposed to estrogen (F_1,9_=5.759, p=0.04, Ηη=0.390) (Figure 6C). This suggests that estrogen exposure at P0 is sufficient to masculinize COX2 expression the cortex.

**Figure 6.**
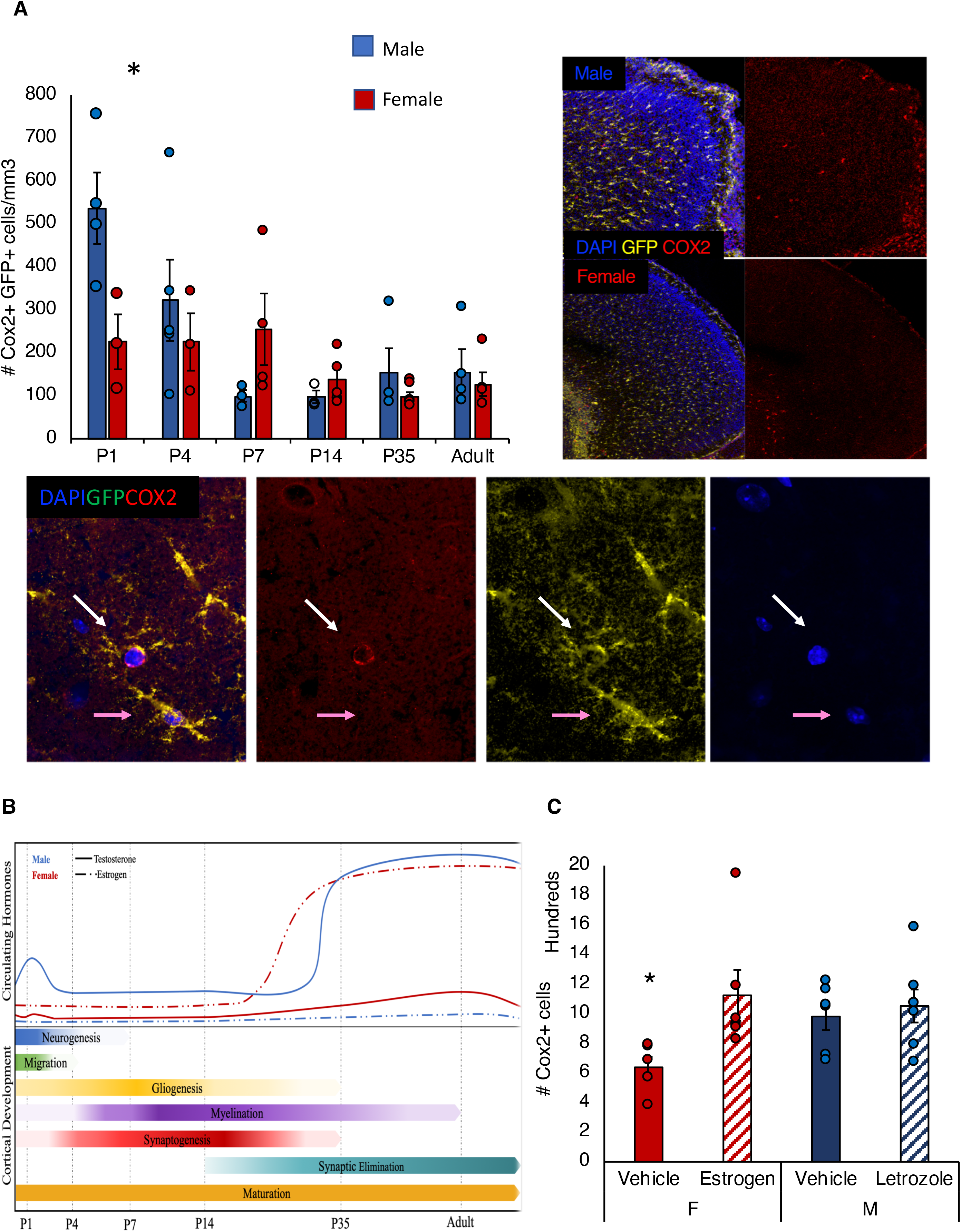
A- Density of COX2+ GFP+ astroglial cells in the cortex across examined timepoints. Bars represent group means, error bars are displayed as ± SEM. Representative mosaic reconstruction of Cox2, eGFP staining in a male and female mouse within the PFC region of the cortex at P1 (top). A 63X+oil image highlighting a Cox2+, eGFP+ astroglia (white arrow) and a Cox2-, eGFP+ astroglia (pink arrow) in the PFC (bottom). B- top panel- Graphical illustration of arbitrary levels circulating gonadal hormones (testosterone, solid lines; estrogen, broken lines) in males (blue) and females (red) overlay with ages examined in the study. B- lower panel- important cortical neurulation events represented by colour saturation. Darker saturation indicates when a process is more active, lower saturation indicates when a process is less active. (Adapted from: Ellis 2004; Gillies and McArthur, 2010; Anderson, 2003; Estes and McAllistter, 2016). C- Density of of COX2+ cells in the cortex at P1 in males and females treated with estradiol or letrozole. Bars represent group means, error bars are displayed as ± SEM. * indicates significance between treatment and vehicle.

### (Sex differences in) Astrocyte differentiation and relevance to disorders

We investigated the relevance of the male and female co-expression network to human disease, by testing the network modules for overlap with disease ontologies (DOs) (see Methods). We focused on the subset of the modules in overlap with astrocyte subtype markers, and report only results for brain-specific disorders (see Supplementary Data File 4 & Supplementary Data File 5, Tab7). In females, top scoring DO terms are intellectual disability and mental deficiency (Module F2); intellectual disability, bipolar disorder, epilepsy, schizophrenia, cognitive delay, autistic disorder, Alzheimer’s disease (Module F3); Unverricht-Lundborg syndrome, lysosomal storage diseases, amyloidosis, progressive myoclonic epilepsies (Module F6); epilepsy seizures, Alzheimer’s disease, schizophrenia, intellectual disability, major depressive disorder, neurodegenerative disorders, autistic disorder, Parkinson’s disease (Module F9). In males, top scoring DO terms are intellectual disability (Module M1); mitochondrial diseases, Alzheimer’s disease, neurodegenerative disorders, epileptic encephalopathy, seizures, bipolar disorder, epilepsy, schizophrenia, autistic disorder, Parkinson’s disease (Module M2); peripheral neuropathy (Module M3); mitochondrial diseases, Leigh disease (Module M6); intellectual disability (Module M8); microcephaly, ciliopathies, intellectual disability (Module M11); epilepsy, bipolar disorder, mental depression, unipolar depression, depressive disorder, Alzheimer’s disease, schizophrenia, Intellectual disability, autistic disorder (Module M12). Notably, many of the diseases are in overlap between males and females, and there does not appear to be disease specificity per se as a wide variety of neurological, psychiatric, and developmental disorders are called. Importantly, however, each of these diseases have been linked to astroglial dysfunction either primarily or in a reactive capacity and therefore the wide variety of diseases called likely speaks to the general involvement of astroglia in all neural processes (Molofsky et al., 2012). Modules associated to differentiated astrocyte subtypes, i.e. increasing with age, are enriched in neurodegenerative disorders terms. Interestingly, these same modules also show enrichment in autism-associated genes, and only modules F9 and M2 show enrichment in Parkinson’s disease genes, unlike Alzheimer’s and schizophrenia. Perhaps reflecting regional involvement in disease pathology, with Parkinson’s origins to mid-brain-striatal functions rather than cortical origins. Module F6, associated with astrocyte progenitor cells, is enriched in epilepsy related genes including a number of GABA transporters and GABA receptor subunits (*gabra3, gabrg2, gabrb2, slc6a1*), unlike its M8 counterpart. Interestingly, both astroglial cells and GABA subunits are central to epilepsy pathology (Coltour and Steinhäuser, 2015; Fritschy, 2009), and sex differences in gaba subunits have been linked to sex differences in epilepsy (Li et al., 2007) however, whether astroglial expression of these subunits is related to developmental sex differences in epilepsy susceptibility is unknown. Finally, the “early peak” module M6 associated with intermediate progenitor cells, is enriched in mitochondrial diseases terms. This male-specific expression is consistent with mitochondrial diseases having a higher incidence in males, thought to be due to uniparental or maternal inheritance (Beekman et al., 2014).

## Discussion

In the current study, we generated a translatome database of cortical astroglial cells across development in both male and female mice. We show two clear astroglial phenotypes across development, an early (before P14) and late (after P14) phenotype. These early and late gene expression profiles reflect astroglia associated with early developmental functions related to gliogenesis and synaptogenesis and astroglia associated with functions involved in maturation, synapse elimination and maintaining a stable, homeostatic environment in adulthood (Figure 7D).

Surprisingly, cortical astroglial cells showed very little sexual dimorphism in either morphology or gene expression particularly in adulthood when very few DEGs were not sex-chromosome associated genes. Importantly, to assess basal differences between males and females in a conservative way, we chose to examine females during the metestrus phase, when ovarian hormones are at their lowest and therefore more similar to levels observed in males. As such, it is likely that greater sex differences in the astroglial translatome would be observed during other estrous cycle phases, such as in proestrus, when estrogen peaks. Indeed, we have previously demonstrated changes in cortical astroglial expression of GFAP and fibroblast growth factor 2 (FGF2), an astroglial mitogen and growth factor across the estrous cycle^49^, and a number of studies have shown functional, morphological and protein expression changes in astroglia in response to sex hormones (Verkhratsky and Nedergaard, 2018; Azcoitia et al., 2010) Together these results would suggest that astroglial cells show phenotypic changes in response to ovarian hormones (or activational effects of hormones), but at least in the neocortex, astroglia do not show basal sexual dimorphism (organizational effects of hormones).

Interestingly, the sex differences in DEGs observed at P7, and P14, were not related to sex-specific canonical pathways or functions per se, but rather, were related to general developmental phenomena such as cell cycle, and neuronal, dendritic and synaptic development, suggesting that astroglia modulate neuronal networks in a sexually dimorphic way. This could, in turn, potentially lead to the formation of sexually dimorphic neuronal networks, and presumably, sexually dimorphic behavioural expression later in life. Given the critical role of the neocortex in psychiatric disease and complex human behaviours^50^, one might imagine that changes in neuronal network development may be induced by estrogen-modulated changes in astroglia. In addition, these may be particularly susceptible to developmental perturbations that modulate hormone expression levels including exposure to stress hormones and environmental endocrine disruptors. Moreover, because hormone levels vary between individuals under normal conditions, this allows for a potential range of sexual differentiation of cortical networks, which would mirror behavioural sex differences observed across species, that are typically expressed on a continuum.

In contrast to the sex differences in gene expression seen in P7-P35, we see few DEGs between astroglia in male and female mice in the early postnatal period and in adulthood. This, in addition to the sex differences in CP of early versus late astroglial phenotype and the apparent early peak of some networks in male mice, suggests that cortical astroglial cells employ different developmental strategies or timing in developmental trajectories to reach a similar adult phenotype, with males accelerating faster than females in early development. The idea that distinct developmental trajectories between the sexes might lead to similar functional results is similar to the concept of compensation put forward by Woolley^51^, which was based on findings of estradiol potentiating glutamatergic synapses, albeit on a much more general level. Earlier DeVries^52^ postulated that sex differences in gene expression patterns over development may actually prevent sex differences in specific regions or ensure that neuronal systems develop and are functional in the presence of cues such as sex hormones that are crucial for stimulating sex differences in target regions. It is possible that the sex differences in gene expression profiles between astroglia ensure that these systems develop “equally” in both sexes to achieve similar functionality in adulthood in the presence of sex chromosomes and hormones that are necessary to differentiate regions critical to sexually dimorphic behaviours, such as the hypothalamus.

The question remains as to what induces sex differences in cortical astroglia gene expression patterns over development. Three potential scenarios emerge: 1) sex chromosome-linked genes directly induce changes in gene expression patterns in astroglia. 2) The perinatal testosterone surge initiates cascades of gene expression patterns in astroglia that modulate cell maturation and network development. 3) Astroglial cells have a similar start and end point in our dataset and it is possible that sex differences in astroglia gene expression represents an active process to compensate for sex differences in developing neuronal imputs to the cortex that arise from dramatic changes in neuronal number, arbours, and synapses on neurons that project from the hypothalamus to the neocortex Importantly, these scenarios need not be mutually exclusive and may be active in parallel systems.

*Scenario 1. Kdm5d* appears as one of the few DEGs between females and males at P1 in our dataset, female expression levels are lower than males. *Kdm5d* is a Y-linked gene that encodes for demethylation enzymes and is required and sufficient to induce sexual differentiation of gene expression^53^. Indeed, changes in gene methylation may have profound effects that persist throughout time and cascades of gene expression may or may not be initiated, altering the responsiveness of the cell, and thus the network as a whole. Moreover, even in the absence of the testes determining *Sry* gene, the Y-chromosome has been shown to exert differential effects on developing peripheral and nervous tissue (reviewed in^54^), lending support to the idea that sex chromosome linked genes induce later sex differences in astroglia across development.

*Scenario 2.* Similarly, our results suggest that the perinatal testosterone surge, and consequentially, estrogen, is involved in the induction of developmental sex differences in astroglia. Our weighted network analysis shows enrichment of genes in several modules associated with responses to sex hormones. We show sex differences in the prostaglandin system (including Cox2) and that neonatal estrogen administration induces changes in cortical COX2 expression. Similar changes in astroglia have been demonstrated in the hypothalamus and are sufficient to produce observable sex differences in neuronal networks and associated behaviour^13^. Thus, the endogenous expression of estrogen early in development may facilitate a cascade of cellular maturation that in-turn will alter subsequent genetic programs. As such, the neuronal network will facilitate different programs as a result of the staggered pattern of maturation and communication.

Finally, *scenario 3* suggests that sex differences in the hypothalamus are necessary for a functional behavioural outcome that is overt and essential for sexual reproduction, first suggested by de Vries^52^. However, there are few overt measures that can be linked to cortical functioning that exhibit similar sexual dimorphisms, although sex differences in risk and resilience for psychiatric disease associated with cortical functioning are well-supported. For example, schizophrenia and ASD are more prominent in the male population^8^ and both show changes in developmental pathophysiology that occur both during the prenatal and postnatal periods^55,56^. There may be a relationship between astroglial neurogenic potential and the terminal phases of neuronal migration and construction and maturation of functional cortical networks that buffers females against developmental anomalies which, if left unchecked, may lead to schizophrenia or ASD morbidity.

In-line with this hypothesis, there are a number of models where estrogen treatment improves recovery or where females are protected in developmental injury models. Chronic postnatal hypoxia in rodents is used as a model of premature birth and induces loss of brain volume, particularly in the cortex and hippocampus, and leads to motor and cognitive impairments later in life^57,58^. The loss of cortical volume in the hypoxic model is less pronounced in females^57^ and interestingly, estradiol treatment at the time of the injury improves white matter damage recovery^59^. It is also known that chronic postnatal hypoxia increases astroglial stem cell capacity^25,60^. This process may be enhanced in females allowing augmented recovery from the aversive effects of hypoxia, but may also speak to the nature of enhanced early postnatal plasticity in females compared to males. It is possible that females experience an enhanced or shifted neurogenic period early postnatally, when we have observed an increase in vimentin+ astroglia, that may translate to increased neurogenic potential of astroglial cells in the cortex. Similarly, while overt network or cellular sex differences may not modulate sex differences in risk for psychiatric disease, sex differences in developmental programs may leave one sex more vulnerable to perturbations during that time. Future studies will be needed to determine the functional significance of sex differences in developmental programs. Nevertheless, our data show a clear developmental pattern in cortical astroglia that differs between males and females, while no apparent basal sex differences in astroglia exist.

## Online Methods

### Animals

54 male and female C57/Bl6-AldHl1-L10-GFP transgenic mice generated from our breeding colony maintained at the Carleton University animal facility within the University of Ottawa. Mice were genotyped as in (Simard et al., 2018) and sex was determined by the presence or absence of tissues in the urogenital tract and confirmed by genotyping for the *Zfy* gene. All animals were group-housed until sacrifice in standard (27cm x 21cm x 14cm), fully transparent polypropylene cages with chew block, bedding, house and *ad libitum* access to standard lab chow (2014 Teklad Global 14% protein®) and water. Animals were raised in the standard environment with no outside manipulation except for standard care and to monitor estrous cycle stage (when applicable). The mice were maintained on a 12-hour light/dark cycle in a temperature controlled (21 degrees) facility. All animal use procedures have been approved by the Carleton University Committee for Animal Care, according to the guidelines set by the Canadian Council for the Use and Care of Animals in Research. Mice were overdosed and perfused for immunohistochemistry or rapidly decapitated for TRAPseq processing each on postnatal day 1* (P1) (36 hours postnatal), P4, P7, P14, P35 and adulthood (defined as 7-9 weeks of age) at the same time of day. Developmental time points and subject numbers are summarized in Supplementary Figure 1.

Organizational hormones study-5 pregnant dams were ordered from Charles River and arrived at the Carleton University animal facility at the University of Ottawa at gestational day 15/16 and given 3 days to acclimatize to the environment. Dams were monitored intermittently overnight starting on gestational day 19 using red light. 31 male and female pups were randomly assigned to control or treatment groups approximately 1 hour following birth and sacrificed 35 hours later. The majority of groups in all experiments were formed with subjects from 3-5 separate litters to control for litter effects.

### Estrous Cycle Monitoring

All adult group female animals were monitored daily after P35 to identify stage of estrous cycle using a saline lubricated swab inserted into the opening of the vagina to collect cells from the vaginal wall. Samples were smeared on a glass microscope slide and examined under a 10x objective using a light microscope (VistaVision^TM^). To understand basal organizational sex differences in astroglia, we used female mice in the phase of the estrous cycle with the lowest circulating ovarian hormones that has the least effect on cortical astroglial morphology and protein expression: metestrus^61^. After the stage of estrous cycle (Metestrus, Diestrus, Proestrus and Estrus) had been established for at least two full cycles, females were sacrificed during the light cycle of the metestrus stage.

### Organizational hormones-injections

Estradiol benzoate (Chem Cruze, sc-205314) was dissolved in peanut oil (0.8g/mL) as per Hisasue et al., 2010 to allow for an injection of 20μg in a 25μL injection volume. Letrozole (Sigma-Aldrich, LG545-10MG) was dissolved in 0.3% hydroxypropyl cellulose PBS (Aldrich, 435007-5G) as per^62^, to allow for an injection of 250mg/kg in a 25μL injection volume, sufficient to block all testosterone conversion based on average pup weight of 5g. Estradiol benzoate and letrozole solutions were prepared using glass materials and stored in a glass amber bottle to prevent absorption and disruption from plastics or ultraviolet exposure.

### Animal Sacrifice

#### Rapid Decapitation for immunohistochemistry and qPCR

Animals in the P1 immunohistochemical group were rapidly decapitated and brains immediately placed in a 4% paraformaldehyde (Fisher Scientific) (PFA) solution at 4 °C for 24h after which the brains were switched to 30% sucrose at 4 °C for 24h. Following this period, brains were flash frozen until slicing. For the organizational hormone experiment, pups underwent rapid decapitation 36h following treatment. Brains were harvested and split along the central fissure. One hemisphere was flash frozen at −80°C for analysis in qPCR and the other hemisphere was fixed as above.

#### Cardiac Perfusion for Immunohistochemical Analysis

All mice were sacrificed for tissue collection at the same time of day during the light cycle. All mice in the immunohistochemical group underwent cardiac perfusion aside from those in group P1 (see above). Animals in P4, P7, P14, P35 and adult groups were given an overdose of 44mg/kg sodium pentobarbital (CDMV Canada), followed by intra-cardiac perfusion upon all spinal reflex cessation. Animals from the P4 and P7 groups underwent manual cardiac perfusion with 1mL of saline solution before receiving 1mL of 4% PFA via syringe. Mice in the P14, P35, and adult groups blood were flushed using 10mL of saline through the left atrium followed by 20mL of 4% PFA to fix the tissue. Brains were extracted and placed into vials containing 4% PFA and stored at 4 °C. Following a 24h period, brains were transferred to a 30% sucrose in PBS (Fisher Scientific) solution and placed at 4 °C.

#### Tissue sectioning

Following sucrose treatment, all brains were then flash frozen at −80 °C until sectioning on a Leica (Leica^TM^ CM1900) cryostat (30μm thick). 15 sets of sagittal sister sections were adhered to electrostatic slides (Fisher Scientific^TM^) in rotating order, each slide therefore contained a full representation of the brain for stereological analysis as per previous studies^60,63^.

### Immunohistochemistry

All immunohistochemical processes took place at room temperature (∼21 °C) as per our previous studies^25,60,64^. One representative slide was taken from every subject and all subjects were processed simultaneously. Brain tissue from all groups were prepared for immunohistochemistry using a 10% horse serum (Gibco^TM^) PBS-T (0.3% TritonX (Fisher Scientific)) pre-block solution for 1 hour before being incubated with the respective primary antibody solution (see Supplementary Figure 1). Primary antibodies were diluted in 10% horse serum PBS-T (0.3% TritonX). Following approximately twenty-four hours of primary antibody incubation, slices were washed in a 1x PBS solution 3 times to remove unbound antibodies before being incubated with the species-appropriate fluorescein conjugated secondary antibody for visualization. Secondary antibodies were diluted in 10% horse serum PBS-T (0.3% TritonX) and incubated for 2 hours before slides were submerged in a 1x PBS wash 3 times to remove unbound antibodies. Slides were then coated with a nuclear stain, DAPI with hard setting mounting medium (VectorLabs), to fix glass cover slips and allowed to set before analysis. See Supplementary Figure 1 for antibodies and dilutions used.

### Stereological Analysis

Unbiased estimates of cell numbers were obtained through use of Zeiss AxioImager M2 with ApoTome motorized fluorescent microscope (Carl Zeiss, Thornwood, NY, USA) in conjunction with a motorized stage and a computer running on Windows 7 using the program StereoInvestigator^TM^ (MicroBrightfield, Colchester, VT, USA). Serial sagittal sections of the right hemisphere obtained through cryosectioning at 30μm on 15 sister sections were used for stereological analysis. Contours encompassing the whole right hemisphere neocortex on each section were drawn as boundaries in StereoInvestigator^TM^. Cells were counted for expression of individual and/or co-expressed proteins using the optical fractionator probe at 40x. Sampling grids were optimized for cortical contours to include at minimum 3 sampling sites per contour to allow for a systematic and unbiased method to estimate cell density and cell quantification for right hemisphere of the cortex regardless of cell shape, size, orientation, spatial distribution, or post-mortem brain shrinkage^65^. Sampling boxes are randomly placed by StereoInvestigator^TM^ within the sampling frame measuring 150μm x 150μm x 30μm with 3 of 6 exclusion borders. Total number of cells per count are reported via StereoInvestigator^TM^ output. For analysis of astroglial morphology, StereoInvestigator^TM^ (MBF, Vermont ^TM^) software on a Zeiss Observer with Apotome (Zeiss^TM^) was employed.

We identified and distinguished between radial and non-radial astroglial cells, counting these using unbiased sampling via the StereoInvestigator (MBF, Vermont ^TM^) at 40x. Representative confocal mosaic images used for photomicrographs were taken using ZEN software (Zeiss^TM^) with the Airyscan 800 microscope (Zeiss^TM^).

### TRAPseq

Translating ribosome affinity purification (TRAP)-sequencing was performed as per^66^. Briefly, eGFP-positive animals were rapidly decapitated and the brains removed and placed into a 1000x activated cyclohexamide (CHX) (100μg/ml, Sigma, dissolved in methanol, American Bioanalytical) dissection buffer at which point the cortex, removing any remaining white matter was dissected and flash-frozen in CHX dissection buffer at −80°C. When appropriate, brains were thawed on ice in homogenization buffer containing protease inhibitors, 1000x activated CHX, and RNAse inhibitors. Tissue samples were mechanically homogenized and then incubated on a rotisserie with anti-GFP (HtzGFP_04 (clone19F7) and HtzGFP_02 (clone 19C8): Memorial Sloan-Kettering Monoclonal Antibody Facility) coated biotinylated magnetic beads (Streptavidin MyOne T1 Dynabeads; Invitrogen # 65601) coated with Protein L (Fisher # PI-29997) at 4 °C overnight. Following incubation with magnetic beads, the samples were collected on a magnet (DynaMag-2; Invitrogen #123-21D) and the unbound fragment was collected and frozen at −80 °C. Bound mRNA fragments washed in polysome buffer underwent RNA extraction with DNAse treatment using Absolutely RNA Nanoprep Kit (Stratagene #400753).

RNA samples were sent to the Yale K.E.C.K facility for sample QC as per Simard et al. 2018: RNA quality was determined using a nanodrop and RNA integrity was determined by running an Agilent Bioanalyzer gel (RINs >8). mRNA was purified from approximately 500ng of total RNA with oligo-dT beads and sheared by incubation at 94C. Following first-strand synthesis with random primers, second strand synthesis was performed with dUTP for generating strand-specific sequencing libraries. The cDNA library was then end-repaired and A-tailed, adapters were ligated and second-strand digestion was performed by Uricil-DNA-Glycosylase. Indexed libraries that met appropriate cut-offs for both were quantified by qRT-PCR using a commercially available kit: Kapa Library Quant Kit (Illumina) (KAPA Biosystems # KK4854-07960298001) and insert size distribution was determined with the LabChip GX or Agilent Bioanalyzer. Only samples with a yield of ≥0.5 ng/ul were used for sequencing.

Sample concentrations were normalized to 10 nM and loaded onto Illumina Rapid or High-output flow cells at a concentration that yields 130-250 million passing filter clusters per lane. Samples were sequenced using 75bp paired-end sequencing on an Illumina HiSeq 2500 according to Illumina protocols. The 6bp index was read during an additional sequencing read that automatically follows the completion of read 1. Data generated during sequencing runs were simultaneously transferred to the YCGA high-performance computing cluster. A positive control (prepared bacteriophage Phi X library) provided by Illumina is spiked into every lane at a concentration of 0.3% to monitor sequencing quality in real time.

### Quantitative Real-Time PCR

To validate the enrichment of astroglial cell genes in our bound, immunoprecipitated fragment, the astroglial specific genes GFAP and Glul expression levels were compared between input and TRAP-immunoprecipitated fragments. In addition, we evaluated RbFox3 and Pvalb gene expression assays to validate a concurrent decrease in neuronal gene expression. 20ng of RNA (concentration determined using NanoDrop (Thermofisher Scientific)) was used to produce cDNA using SuperScript IV First Strand Sythesis System (ThermoFisher Scientific). Best-coverage TAQMAN assays (Life Technologies) were run as per manufacturer’s instructions and analyzed using the Applied Biosystems 7500 Real-Time PCR system and software.

To validate the TRAP-seq results we used RNA from all bound (*n*=55), immunoprecipitated fragments to produce cDNA using SuperScript III First-Strand Synthesis SuperMix (Thermo Fisher Scientific; catalog #11752050). We then performed a preamplification reaction (14 cycles) using TaqMan PreAmp Master Mix (Thermo Fisher Scientific; catalog #4384267), with a pooled-assay mix generated according to user guidelines. Finally, best-coverage TaqMan Gene Expression Assays (see Supp. Figure 3) were run as per the manufacturer’s instructions using the Bio Rad CFX Connect Real Time system and software.

### Statistical Analysis

All immunohistochemical and qPCR data were analyzed for using a 2 (male vs. female) x 6 (P1 vs. P4 vs. P7 vs. P14 vs. P35 vs. Adult) between-subject factorial analyses of variance (ANOVA) design using IBM SPSS Statistics (Version 20.0). Because the treatment differed in males and females, for the organizational hormones study we employed a one-way ANOVA to determine overall significance. Post-hoc analysis using Bonferroni correction of pairwise comparisons for non-orthogonal comparisons were conducted when p<0.05. We re-ran all of the statistics including litter composition (male vs. female) as a covariate and found no significant effects of litter on any of the variables measured (data not shown).

#### RNAseq data analysis

Sequencing reads were mapped to the mouse genome (GRCm38.p5) and the Ensembl (releas-84) transcriptome annotation, using HISAT2 (Pertea et al., 2016). The mapping rates ranged from 74% to 95%. This resulted in a number of mapped pairs from ∼19.5 million up to 40 million (see Supplementary Table 6 for more details). The resulting SAM files were converted to BAM formant using SAMtools^67^. The gene expression levels as counts were estimated using featureCounts^68^ with the following options: -t exon; -g gene_id; -F “GTF”; -B; -s 0; -p; -a. Counts from technical replicates were added. Raw read counts were then filtered requiring more than 1 CPM (counts per million) in at least 25% of the samples and 16208 survived the filter and were considered for further analysis. We next used the sva Bioconductor package (Leek et al., 2012) to identify and correct for latent variables. Hierarchical clustering (using the genes with a sd in expression level above the 95%) pre- and post sva correction show the sample-to-sample relationship and relative clustering (see Supplemental Figure 2). FastQC v0.10.1 and RNA-SeQC (v1.1.8) were used for QC (see Supplemental Table QC). The edgeR function rpkm was used to estimate gene expression levels as RPKM.

#### Differential expression analysis

Differential expressed genes were then inferred using the edgeR pipeline^69^, using the trended dispersion to estimate the biological variance and the GLM capability of the package. Nominal p-values from differential expression analysis were FDR corrected, and an FDR cut-off of 0.05 was used for all the tests.

#### Functional enrichment analysis

ConsensusPathDB^70^ was used to test differentially expressed genes for overrepresentation in Gene Ontologies and Canonical Pathways.

#### Weighted gene co-expression network analysis

We used Weighted Gene Co-expression Network Analysis (WGCNA)^71^ for co-expression network analysis using gene expression estimates (as log2(RPKM+1)), separately, from the 25 male samples and the 26 female samples, after cqn and sva correction. We estimated male and female co-expression networks and modules using the function blockwiseModule, using the bicorr as correlation estimate, a “signed” network type and a minModuleSize=50. The power cut-off was set to 16 for both sets. The analysis produced a male network of 13 modules and a female network of 15 modules, including the grey module of unassigned genes (See WGCNA Summary Data).

#### Transcription Factor Analysis

We used the ChEA3 online suite (Keenan et al., 2019) to test each network module for enrichment in Transcription Factors (TFs), retaining only significant calls at FDR<0.05. We considered all the available databases within ChEA3. This resulted in duplicated significant TFs, which were combined, considering the union of all the reported target genes. We estimate the activity for each TF using the camera gene set enrichment analysis algorithm (Wu and Smyth, 2012). considering the P4 vs P1 difference. FDR corrected for multiple comparisons and used FDR<0.05 as cut-off. Any TF estimated as non-significantly active was not retained for further analysis. We then built a directed network from the remaining TFs, using the known TF-target interaction and the WGCNA adjacency as edge weights.

## Acknowledgements

We would like to thank our funding providers, the Carleton University Discovery Grant to N.S., Natural Sciences and Engineering Research Council of Canada Discovery Grant to N.S., the Canadian Foundation for Innovation to N.S., and the Canadian Research Chairs to N.S..

We thank Dr. Julianna Tomlinson, Dr. Michael Schlossmacher and his laboratory members at the University of Ottawa for providing us with infrastructural support and more. Finally, we would like to thank Dr. Alfonso Abizaid for his invaluable input into the planning and preparation of this research project.

---

**Supplemental Figure 1.**
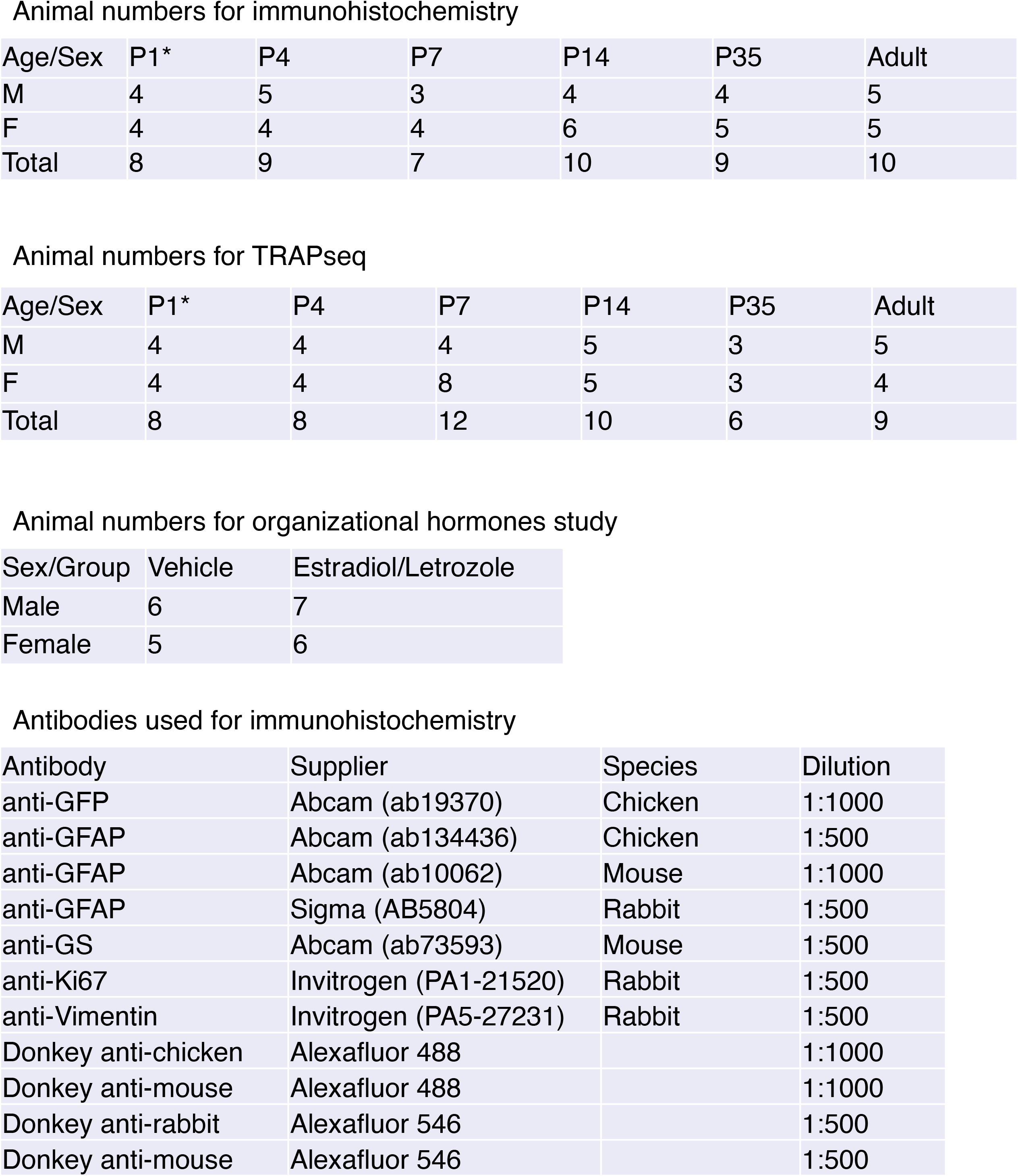

**Supplemental Figure 2.**
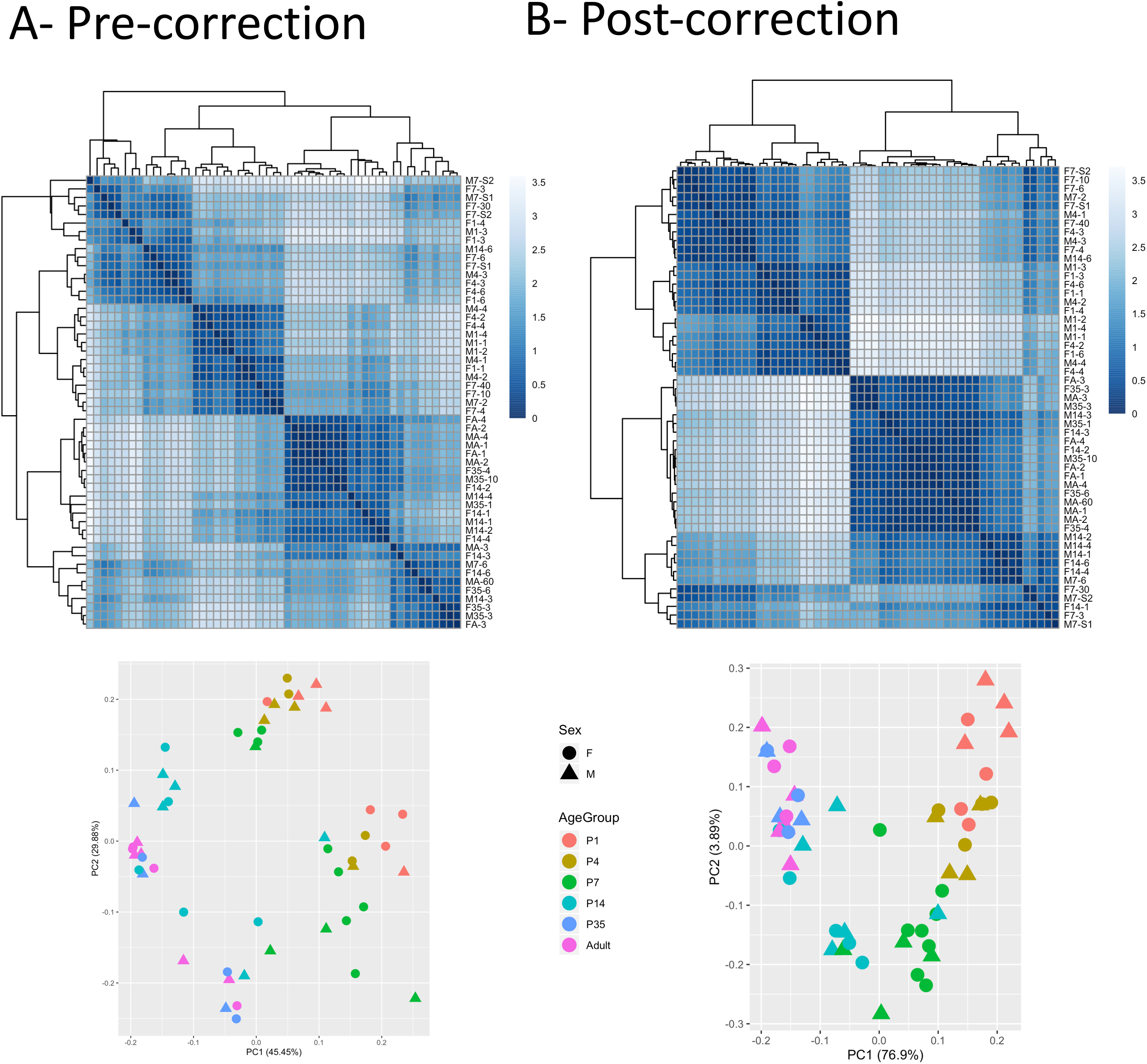
Top- Hierarchical clustering (using the genes with a sd in expression level above the 95%) pre- and post sva correction show the sample-to-sample relationship and relative clustering Bottom- Principal component analysis plots. Represented are the sample distribution along the first principal component (PC1) and then second principal component (PC2). Shows in brackets os the percentage of the variance captured by PC1 and PC2. Circles: female samples; squares: male samples. Symbols are color coded by age.

**Supplemental Figure 3.**
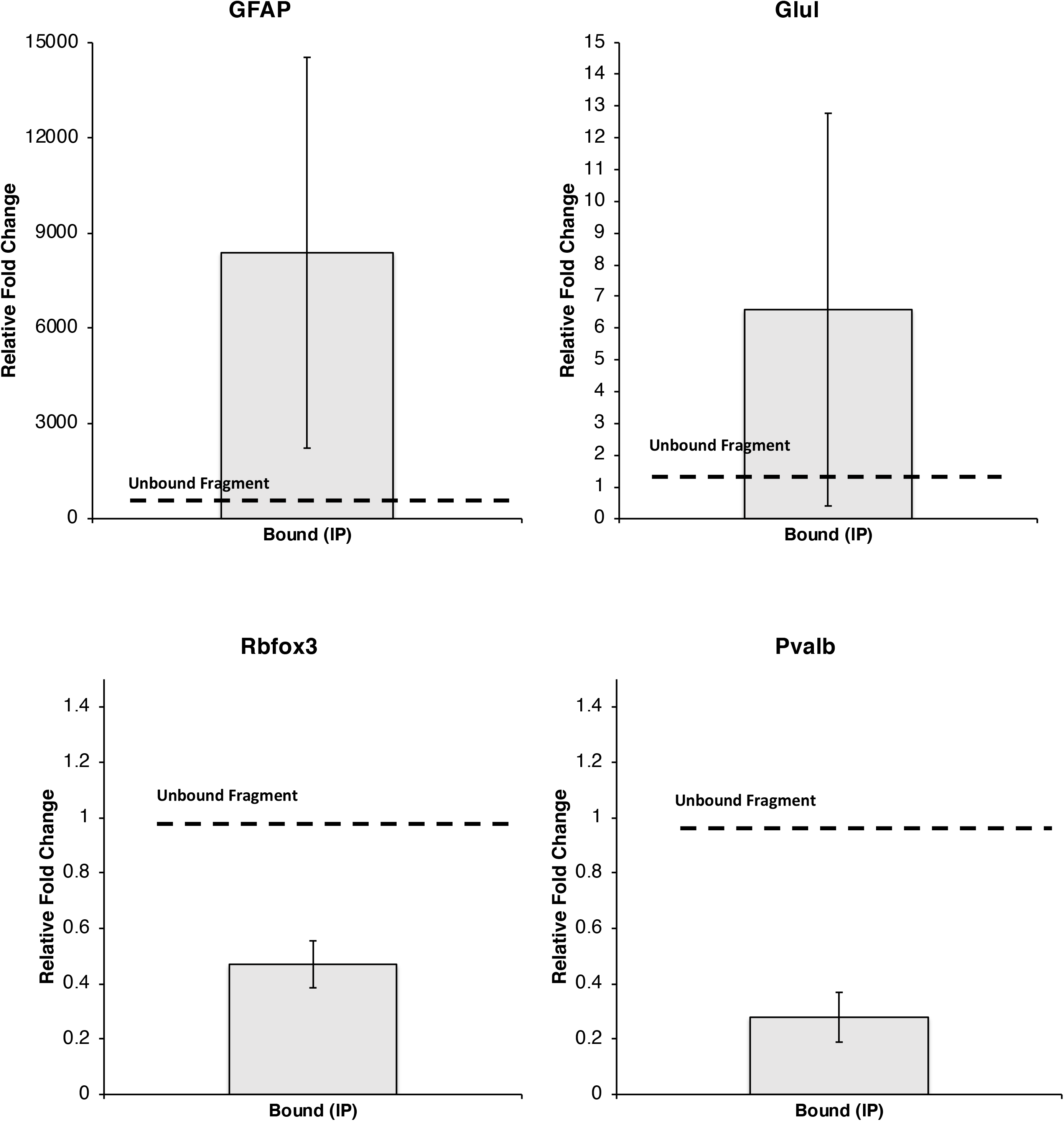

**Supplemental Figure 4.**
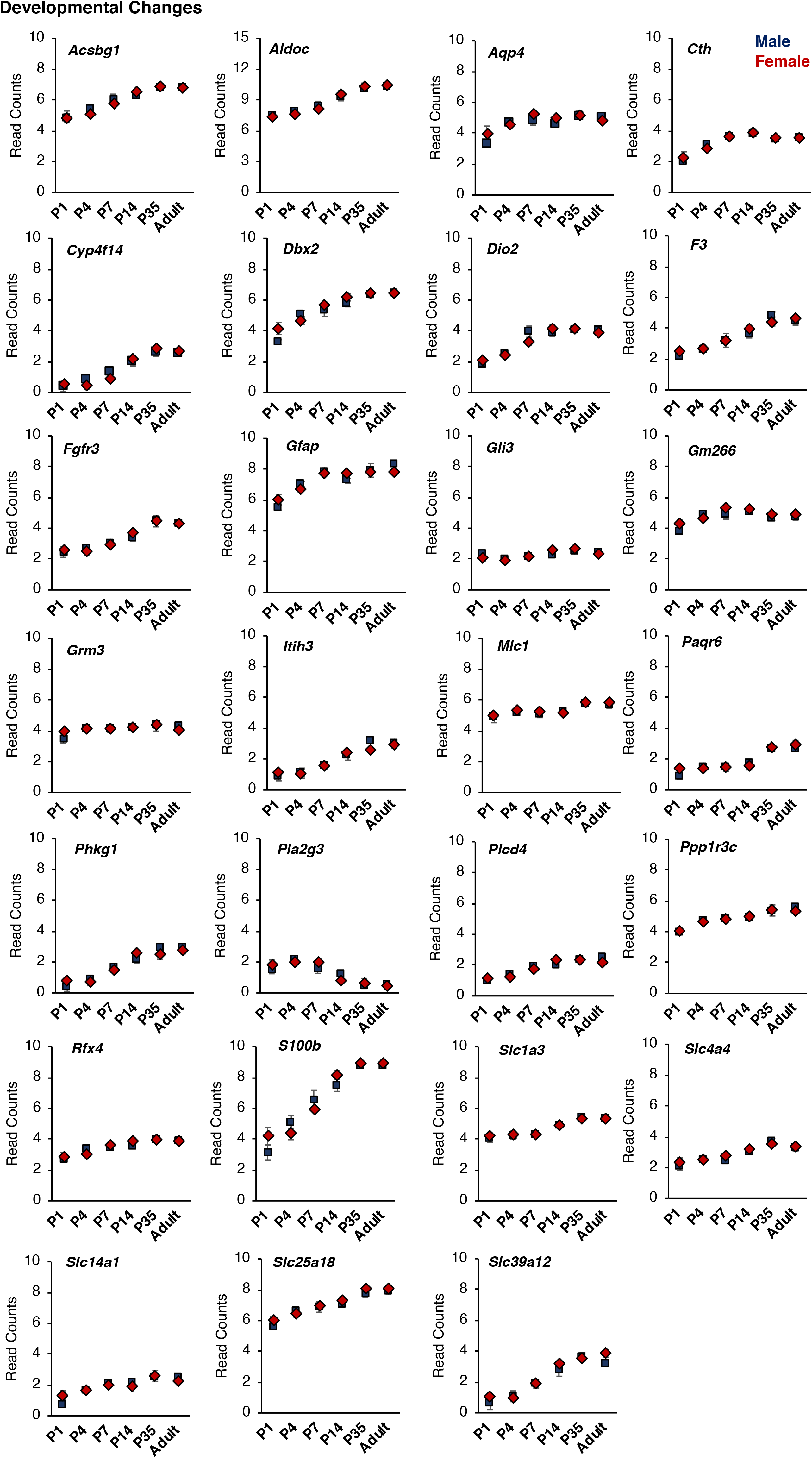

**Supplemental Figure 5.**
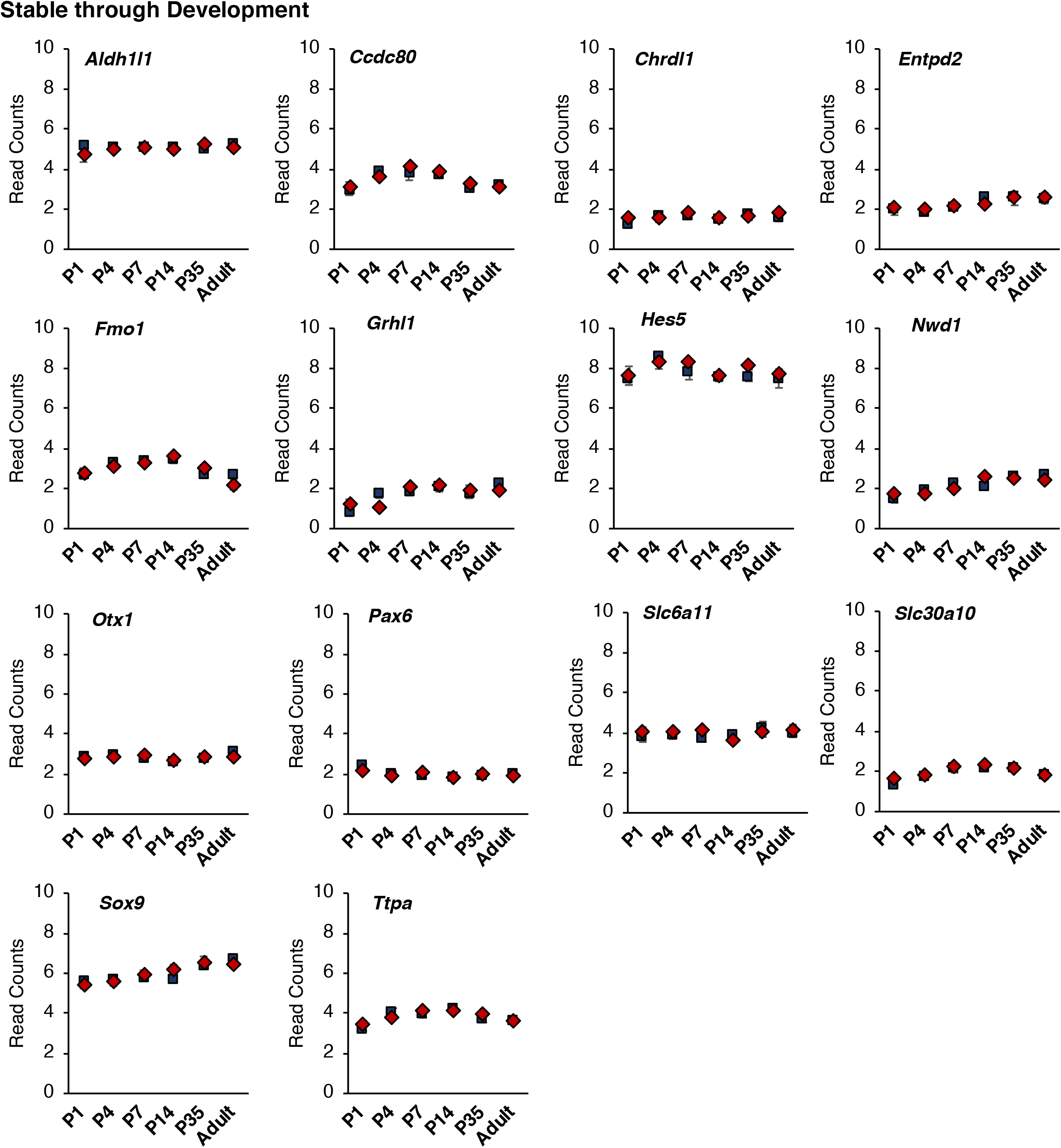

**Supplemental Figure 6.**
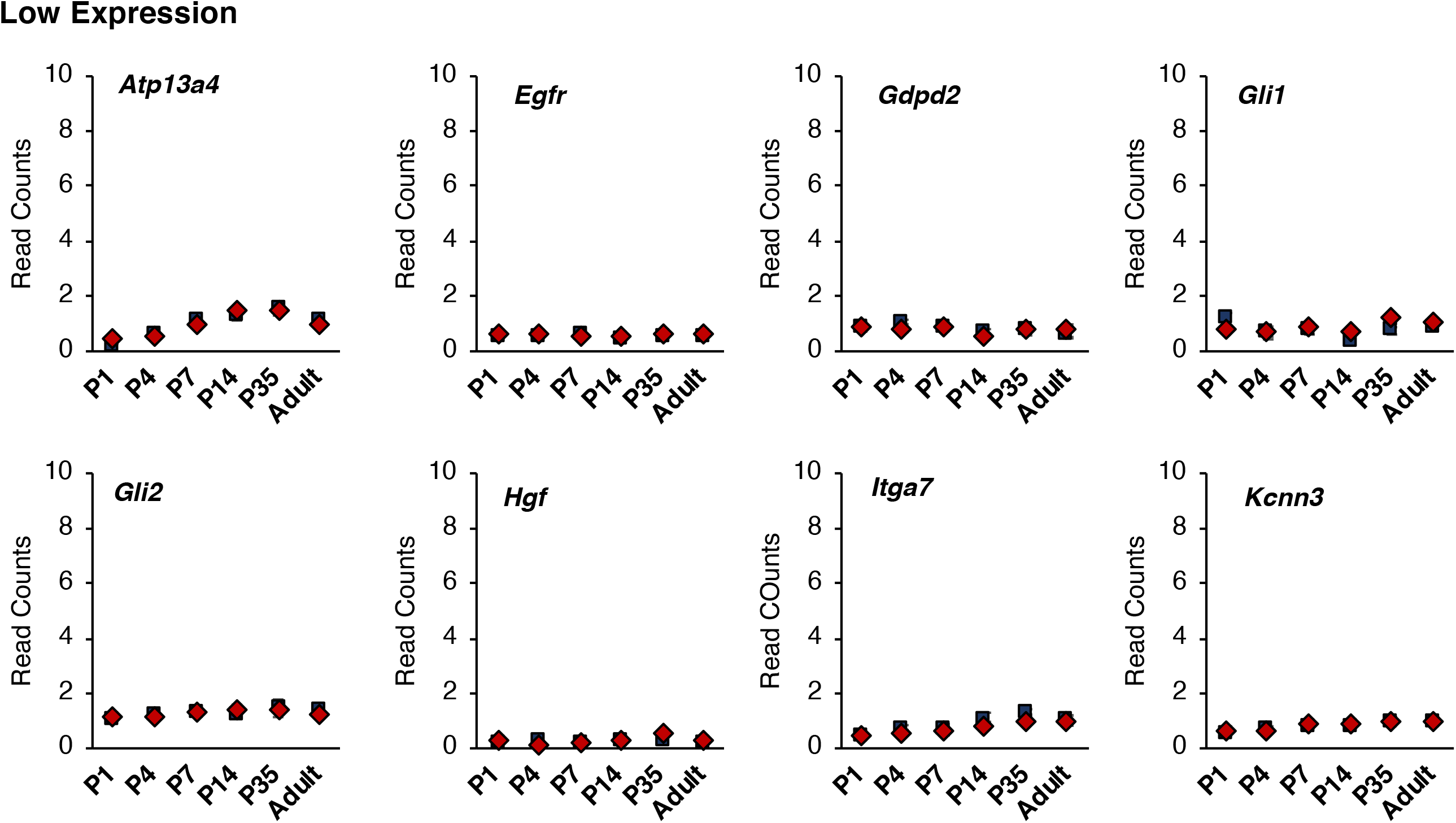

**Supplemental Figure 7.**
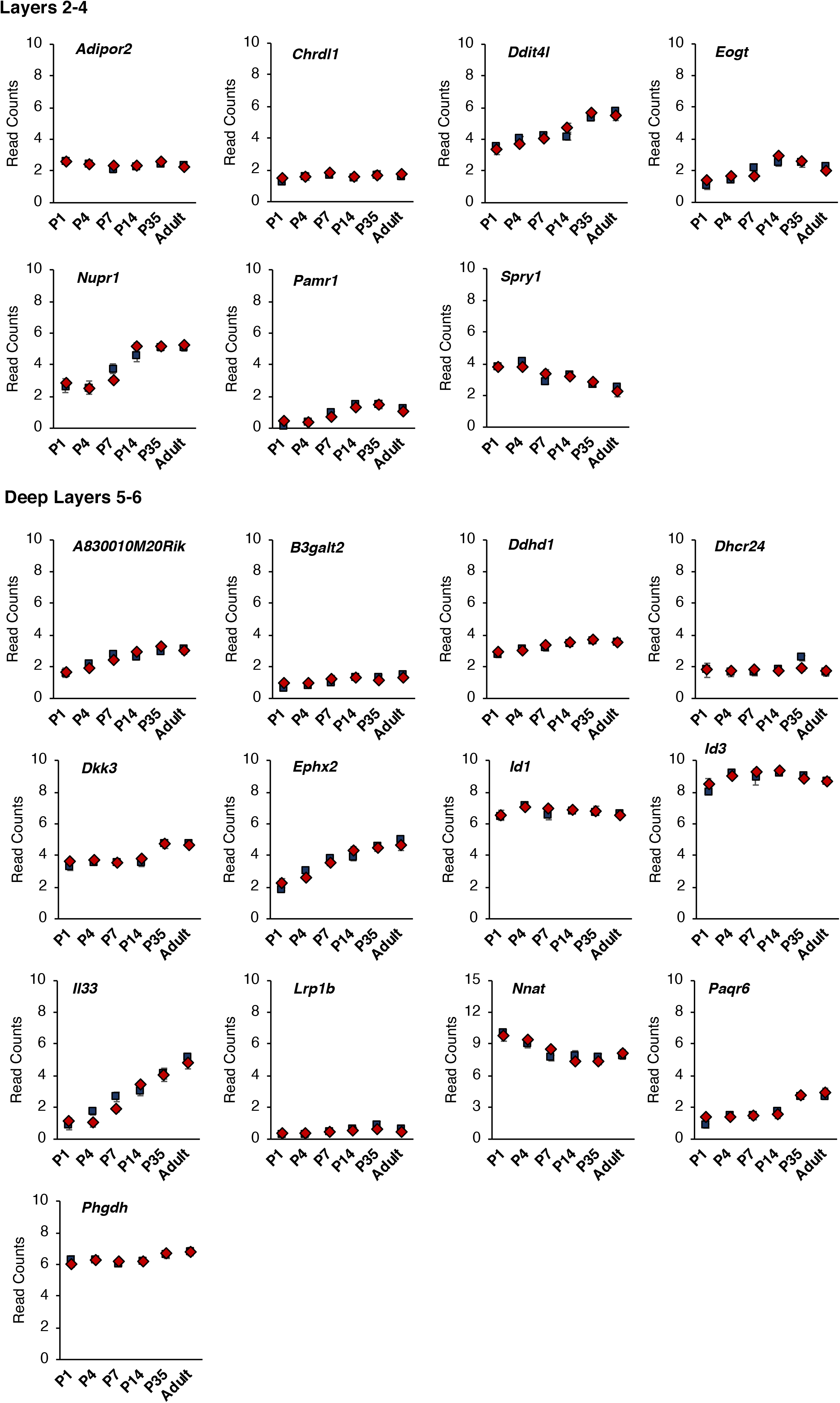

**Supplemental Figure 8.**
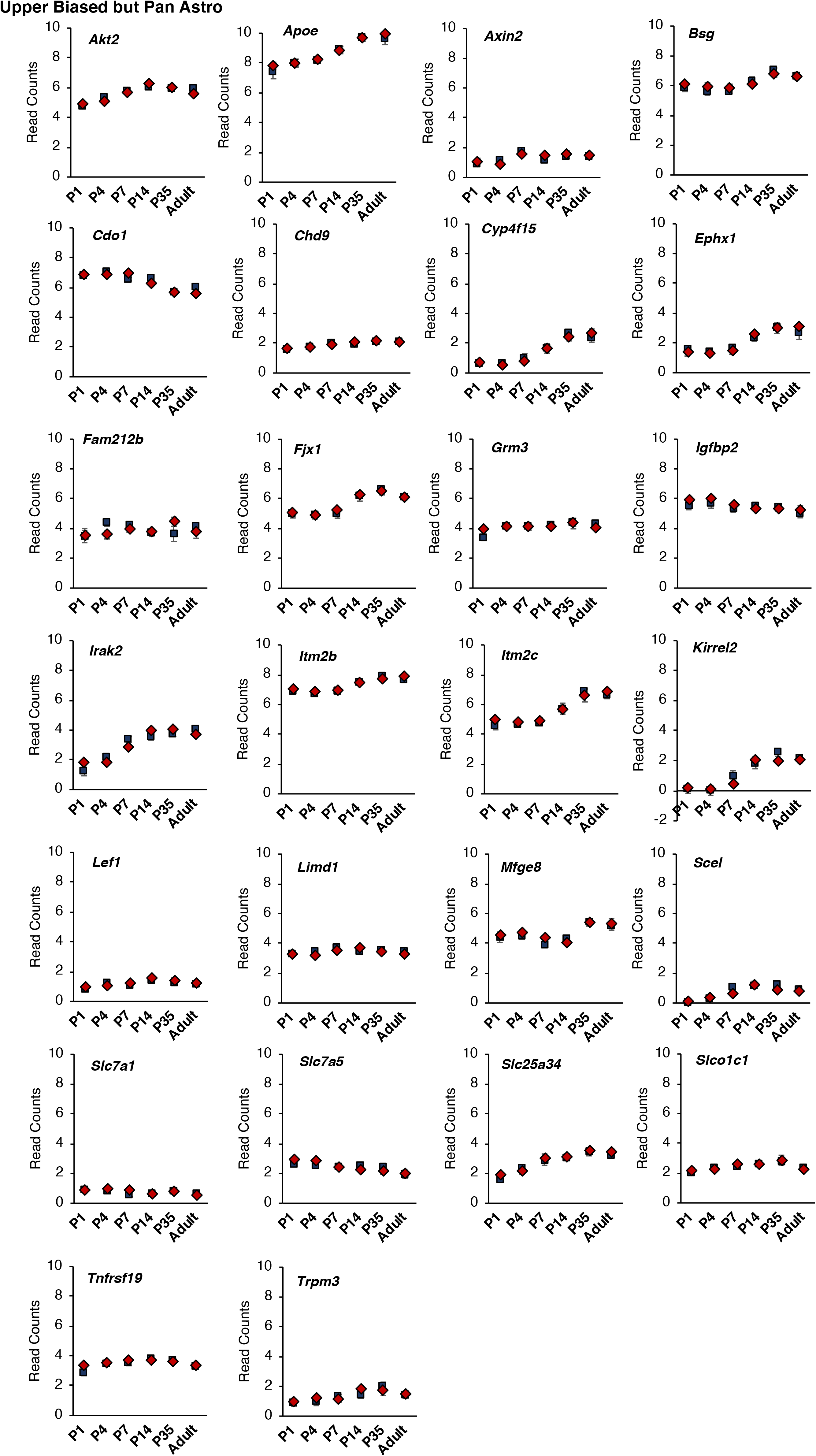

**Supplemental Figure 9.**
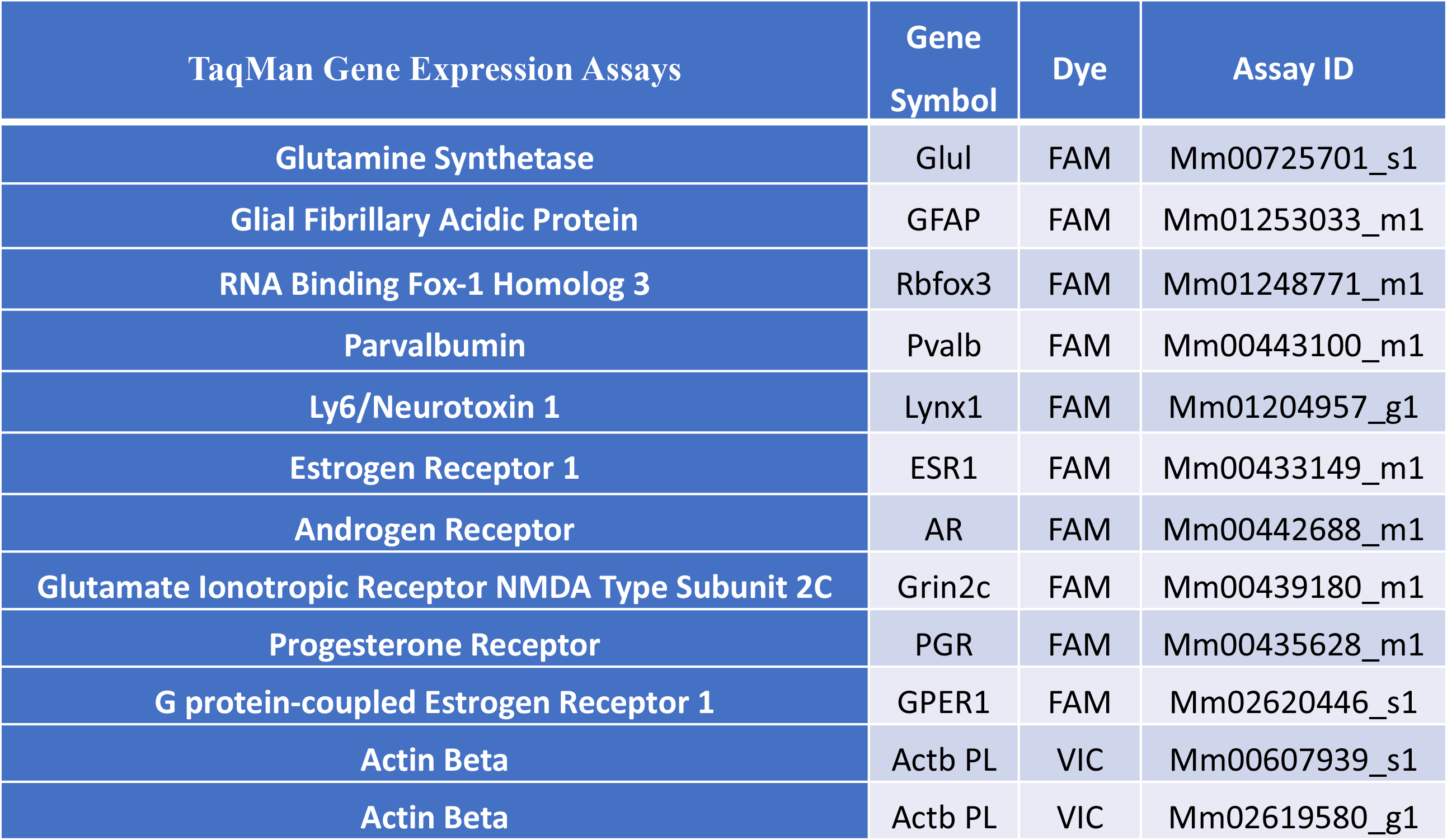

**Supplemental Figure 10.**
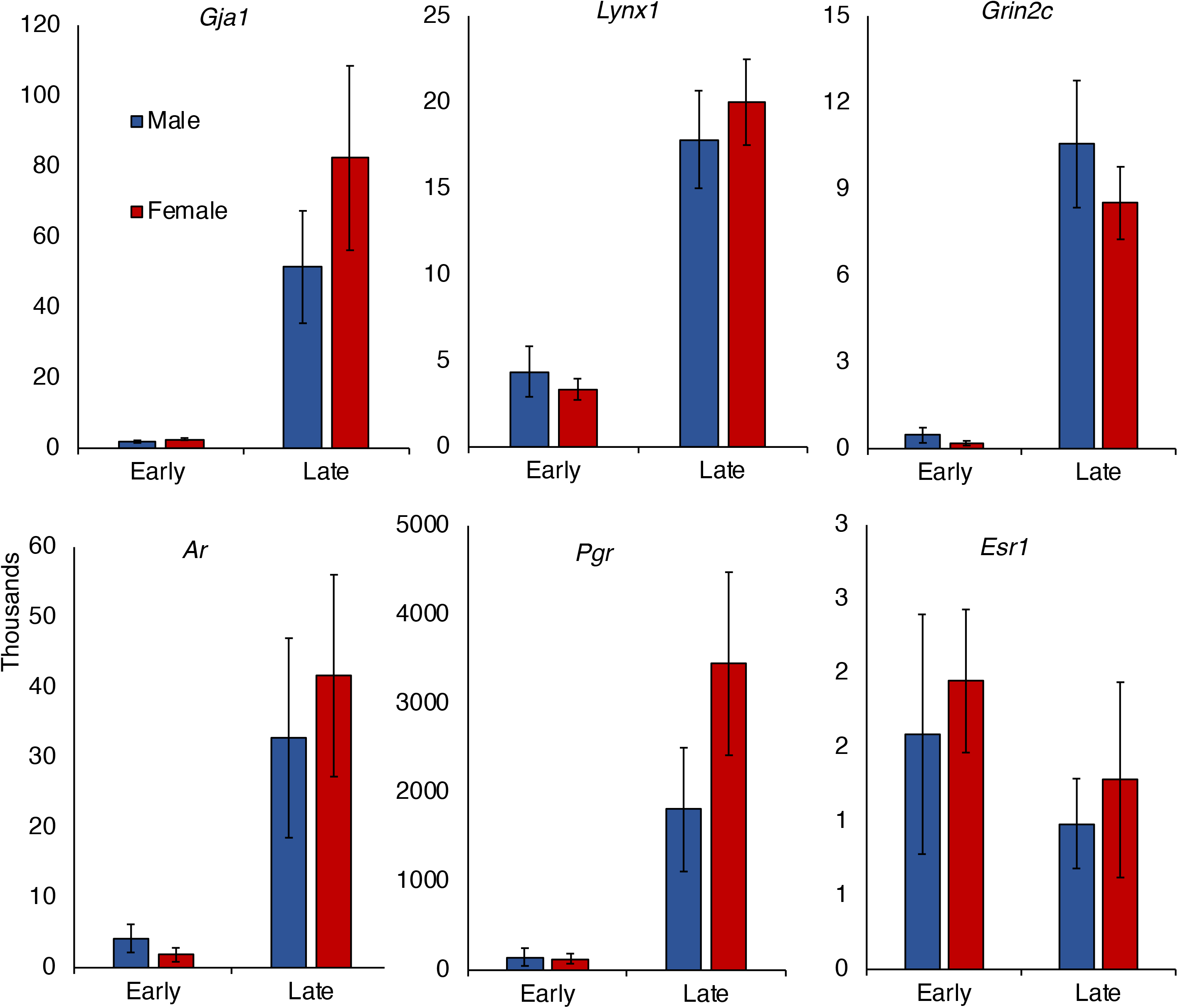
Q-RT-PCR validation of gene expression levels between early (P1-P7) and late timepoints (P14-Adult). Expressed as fold change.

**Supplemental Figure 11.**
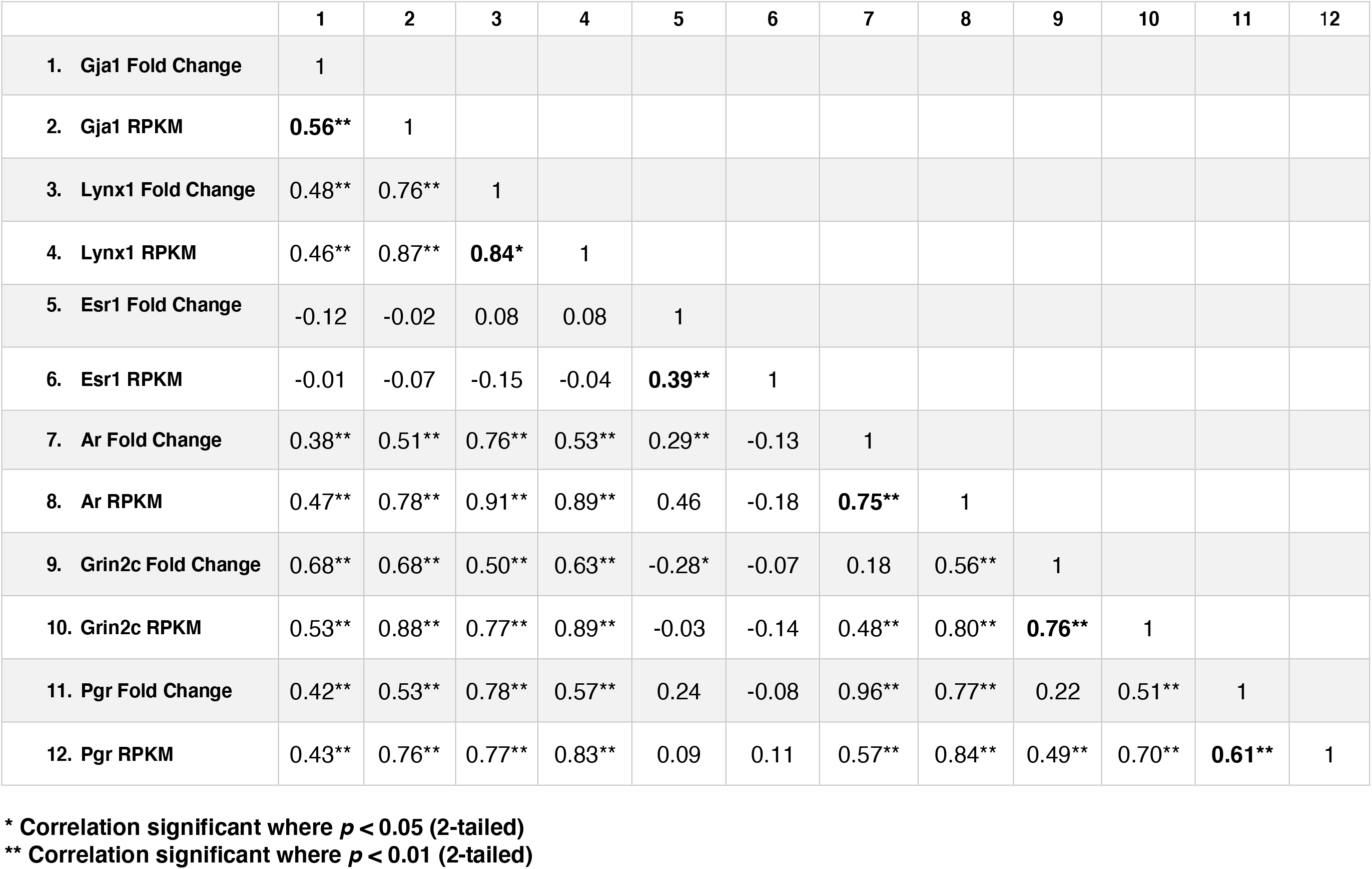
Correlation Table. Pearson correlations between Q-RT-PCR (fold change) validation of gene expression levels in TRAP samples and RPKM levels from TRAP-RNASeq.

**Supplemental Figure 12.**
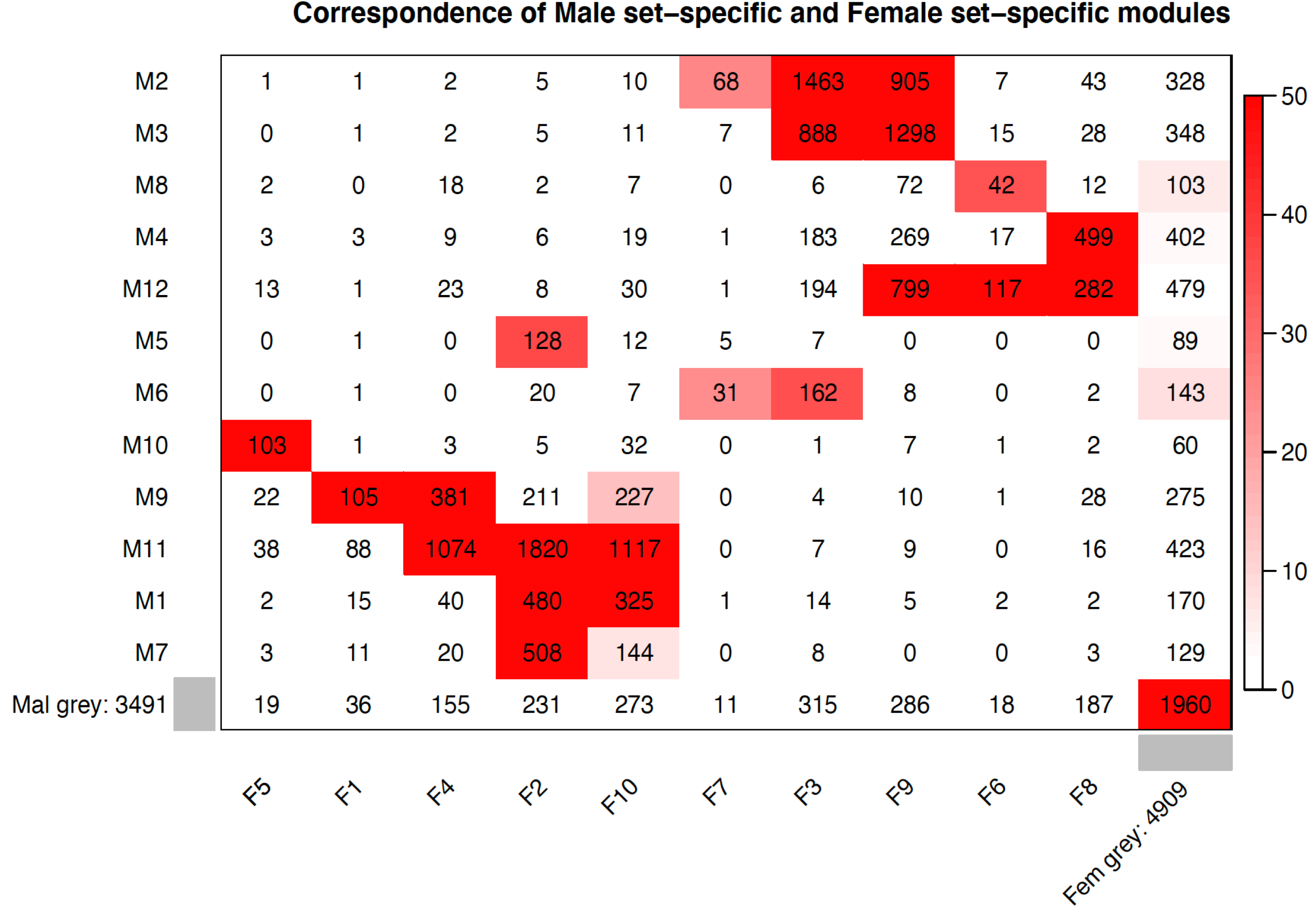
Contingency table of the male modules (rows) and female modules (columns). The module color, and the total number of genes in the module label both the rows and the columns. Numbers within the table represent the counts of genes in the intersection of the corresponding row and column module. The table is color-coded by the −log_10_ of the Bonferroni corrected Fisher exact test p value (legend on then right).

